# Dynamic categorization rules alter representations in human visual cortex

**DOI:** 10.1101/2023.09.11.557257

**Authors:** Margaret M. Henderson, John T. Serences, Nuttida Rungratsameetaweemana

## Abstract

Everyday perceptual tasks require sensory stimuli to be dynamically encoded and analyzed according to changing behavioral goals. For example, when searching for an apple at the supermarket, one might first find the Granny Smith apples by separating all visible apples into the categories “green” and “non-green”. However, suddenly remembering that your family actually likes Fuji apples would necessitate reconfiguring the boundary to separate “red” from “red-yellow” objects. This flexible processing enables identical sensory stimuli to elicit varied behaviors based on the current task context. While this phenomenon is ubiquitous in nature, little is known about the neural mechanisms that underlie such flexible computation. Traditionally, sensory regions have been viewed as mainly devoted to processing inputs, with limited involvement in adapting to varying task contexts. However, from the standpoint of efficient computation, it is plausible that sensory regions integrate inputs with current task goals, facilitating more effective information relay to higher-level cortical areas. Here we test this possibility by asking human participants to visually categorize novel shape stimuli based on different linear and non-linear boundaries. Using fMRI and multivariate analyses of retinotopically-defined visual areas, we found that shape representations in visual cortex became more distinct across relevant decision boundaries in a context-dependent manner, with the largest changes in discriminability observed for stimuli near the decision boundary. Importantly, these context-driven modulations were associated with improved categorization performance. Together, these findings demonstrate that codes in visual cortex are adaptively modulated to optimize object separability based on currently relevant decision boundaries.

## Introduction

Perceptual categorization is a fundamental cognitive ability that allows us to organize and understand the myriad stimuli encountered in our sensory environment. By forming categories, observers are able to generalize existing knowledge to new incoming inputs, facilitating efficient perception and decision-making (Bruner, 1957; Freedman & Assad, 2016). Within the visual system, categories can capture divisions within the natural structure of a stimulus space (Rosch et al., 1976) or can reflect the learning of arbitrary discrete boundaries along stimulus dimensions that would otherwise be represented continuously (Ashby & Maddox, 2005). At the same time, categorization in the real world is a highly dynamic cognitive process, in which the category membership of stimuli may change over time. For example, when making a categorical decision about produce at the farmer’s market, depending on our goals we might think of carrots in the same category as lettuce (vegetables) or the same category as tangerines (orange colored items). Perceptual categorization is thus also tightly connected with flexible prioritization of information based on current task demands (Biederman et al., 1973; McAdams & Maunsell, 1999; Desimone & Duncan, 1995). Within contexts where task goals change dynamically over time, the neural mechanisms supporting categorization of sensory stimuli are not yet understood.

Past work has provided some insight into how category learning impacts representations of sensory stimuli. Behaviorally, learning to categorize stimuli in a continuous feature space can lead to perceptual changes such as an increase in sensitivity to changes along a relevant stimulus dimension, and an increase in perceptual discriminability of stimuli belonging to different categories (Goldstone, 1994; Livingston et al., 1998; Newell & Bülthoff, 2002). Such changes are also reflected in the brain – electrophysiology studies in macaques have demonstrated that after learning of a categorization task, neurons in inferotemporal cortex (ITC) become more strongly selective for diagnostic dimensions of stimuli (Sigala & Logothetis, 2002), and neural populations in ITC also contain information encoding the learned category status of stimuli (Meyers et al., 2008; Tanaka, 1996). In human functional magnetic resonance imaging (fMRI) studies, learning to discriminate object categories has been shown to increase neural responses to objects in extrastriate cortex (Gauthier et al., 2000; Op de Beeck et al., 2006) and lead to sharpening of visual representations as measured with fMRI adaptation (Folstein et al., 2015; Folstein et al., 2013; Jiang et al., 2007). Moreover, recent work has shown that learning a decision boundary can alter representations of orientation in early visual areas, with representations becoming biased away from the decision boundary (Ester et al., 2020). At the same time, other work has suggested that the effects of category status on sensory representations are more prominent in prefrontal cortex (PFC) than visual areas. This suggests that the primary role of visual areas may be restricted to perceptual analysis, rather than decision-related processing (Freedman et al., 2003; McKee et al., 2014; Meyers et al., 2008).

From an efficient processing perspective, it is plausible that visual areas play a more active role in decision-making, potentially encoding decision-related variables, task contexts, choices, or motor outcomes. Such coding would enable visual areas to process sensory inputs in a manner conducive to downstream readout. Emerging evidence from rodent studies supports this view. For instance, activity that was thought to reflect random fluctuations in neural representations within sensory areas has been linked to choice-related motor activities and decision outcomes (Musall et al., 2019; Stringer et al., 2019). Furthermore, recent findings indicate that early sensory areas robustly encode task context variables, such as expectations and decision rules, during dynamic decision-making tasks (Ebrahimi et al., 2022; Findling et al., 2023). Yet, the extent to which human sensory areas similarly code for task-related variables and adapt their representations based on contextual changes is unclear.

In addition, the mechanisms by which categorical decision-making flexibly shapes neural representations, particularly in tasks necessitating the switching between distinct decision rules, are not well understood. Prior work has demonstrated that neural populations in PFC can dynamically encode different boundaries depending on the currently relevant task rule (Cromer et al., 2010; Roy et al., 2010), providing one potential neural mechanism for dynamic decision-making. Similarly, a human neuroimaging study using novel objects suggested that representations in frontoparietal areas can encode different category distinctions between objects depending on their task relevance (Jackson et al., 2017). This study also found evidence for similar (albeit weaker) effects in the lateral occipital complex (LOC), suggesting that representations in visual areas may also be modified by task-relevance. Thus, it remains an open question whether and how varying task contexts interact with representations in visual cortex, as well as how these modulations may contribute to downstream task performance.

Here we address these gaps by investigating how neural responses in human visual cortex flexibly adapt to dynamic task contexts, as induced by varying categorization rules. We hypothesized that task context modulates sensory representations such that changes in the decision boundary are actively integrated during the early analysis of sensory information. To examine the effects of categorization within an abstract stimulus space, we generated a two-dimensional space of shape stimuli (Op de Beeck et al., 2001; Zahn & Roskies, 1972) that were viewed by human participants undergoing fMRI scanning. Participants categorized shapes according to different rules: linear boundaries (*Linear-1* and *Linear-2* tasks) or a non-linear boundary (*Nonlinear* task). These task contexts were interleaved across scanning runs, necessitating real-time cognitive adaptation to distinct categorization requirements applied to physically identical stimuli. Each task incorporated both “easy” and “hard” trials drawn from distinct locations in the shape space, enabling us to concurrently examine the influence of perceptual difficulty on decision processes. Using multivariate decoding methods in retinotopically-defined visual areas, we measured shape representations in each categorization task and examined how representations differed across task contexts. We predicted that shape representations would be more discriminable across a given decision boundary when that boundary was relevant for the current task. Findings from our neural data are in line with this account. Importantly, we further show that an increase in neural discriminability is linked to improved task performance.

## Results

We trained 10 human participants to perform a shape categorization task while in the fMRI scanner, with each subject participating in 3 scanning sessions that each lasted 2 hours (Figure 1A). Shape stimuli varied parametrically along two independent axes, generating a two-dimensional shape space, and each condition of the task required shapes to be categorized according to either a linear boundary (*Linear-1* and *Linear-2* tasks) or a nonlinear boundary that required grouping together of non-adjacent quadrants (*Nonlinear* task). These different categorization tasks were performed during different scanning runs within each session, meaning that participants needed to flexibly apply different decision rules depending on the task condition for the current run (see *Methods*). Each task included a mixture of “easy” trials and “hard” trials. On the “easy” trials, a common set of 16 shapes, making up a 4×4 grid which we refer to as the main grid (black dots in Figure 1B), were shown in all tasks, while on “hard” trials, shapes were sampled from portions of the shape space near the active boundary, which made the current task more challenging (light gray dots in Figure 1B).

**Figure 1.**
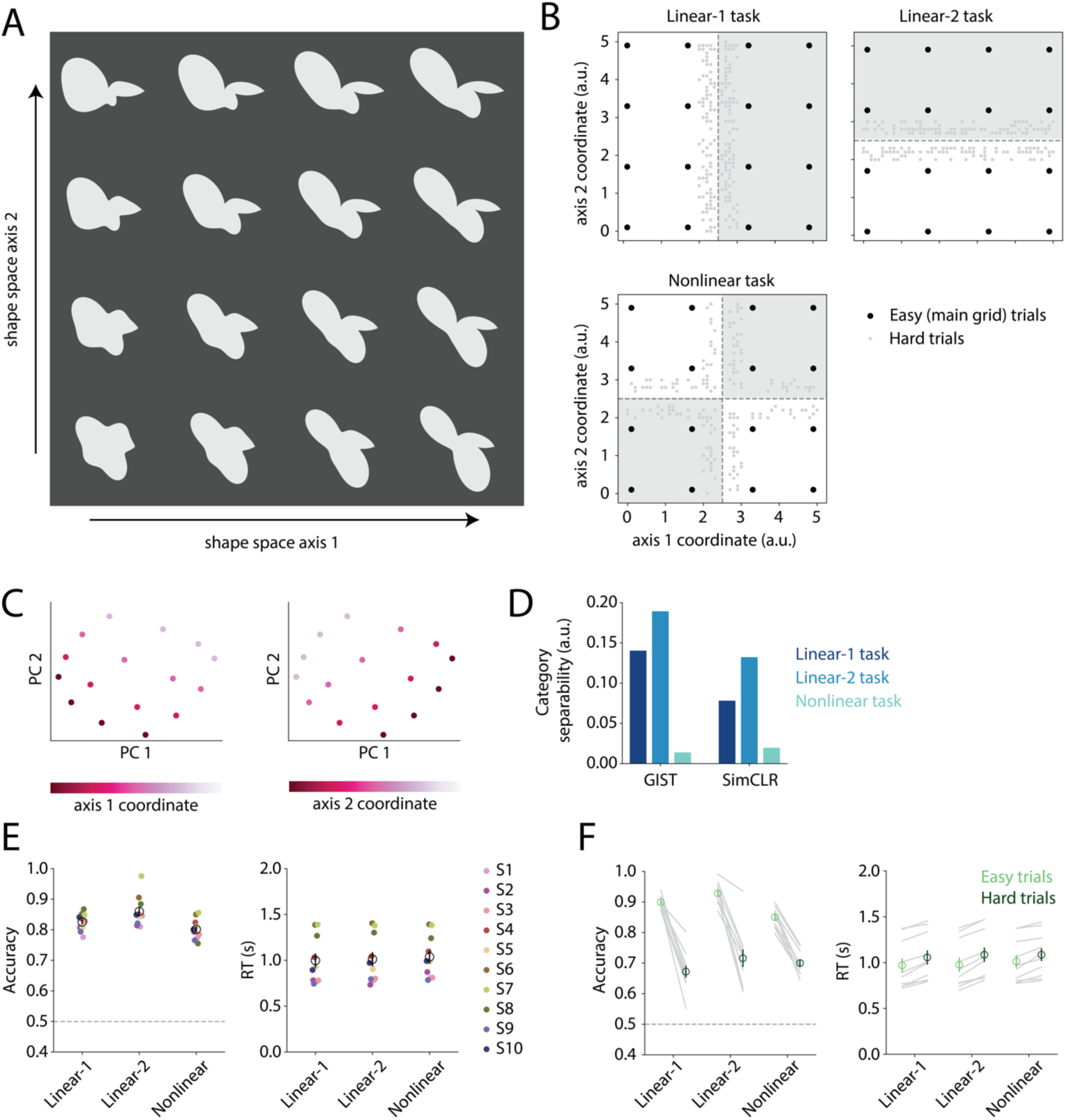
Stimulus set, task design, and behavioral performance. **(A)** Two-dimensional shape space used for categorization tasks in this experiment. Shapes are generated using radial frequency contours (Op de Beeck et al., 2001; Zahn & Roskies, 1972) that vary along two independent dimensions, referred to as axis 1 and axis 2. See *Methods* for more details. **(B)** Illustration of the tasks (*Linear-1, Linear-2, Nonlinear*) performed by participants while in the fMRI scanner. Points in each plot indicate the positions in shape space that were sampled, and dotted lines indicate the relevant categorization boundaries for each task. Black dots represent the 16 positions in the “main grid”, which were sampled on “easy” trials in every task, while light gray dots represent positions that were sampled on “hard” trials, which differed depending on the task. Hard trial shape positions were sampled from the region nearest the relevant categorization boundary. Different tasks were performed during different scan runs. In each task, every trial consisted of the presentation of a single shape (1s), and participants were instructed to respond with a button press indicating which category the presented shape fell into. See *Methods* for more details on task design. **(C-D)** Image similarity analysis: we computed activations from two computer vision models, GIST (Oliva & Torralba, 2001) and SimCLR (T. Chen et al., 2020) for each of the 16 main grid shape images. **(C)** Visualization of a principal components analysis (PCA) performed on the GIST model features, where each plotted point represents one shape in PC space, colored according to the coordinate value along axis 1 (left) or axis 2 (right). **(D)** Quantification of the separability of shape categories within each feature space, computed based on the ratio of between-category to within-category Euclidean distance values. See *Methods* for more details. **(E)** Behavioral accuracy (left) and response time (RT; right) in each task. Dots in different colors represent individual participants; open circles and error bars represent the mean ± SEM across 10 participants. **(F)** Accuracy (left) and RT (right) for each task separated into “easy” and “hard” trials, where easy refers to trials sampling the 16 shapes in the main grid (black dots in B), and hard refers to trials sampling more challenging portions of the shape space for each task (light gray dots in B). Gray lines represent individual participants, open circles and error bars represent the mean ± SEM across 10 participants.

To verify the two-dimensional structure of our shape space, we used an image similarity analysis based on GIST features (Oliva & Torralba, 2001; see *Methods*) to assess the perceptual similarity between shape stimuli. As expected, a principal components analysis (PCA) performed on the GIST features revealed a two-dimensional grid structure, with the two shape space axes oriented roughly orthogonal to one another in PC space (Figure 1C). In addition, measuring the linear separability (based on between-category versus within-category Euclidean distances; see *Methods*) of shapes across each category boundary based on GIST features revealed that shapes were most separable across the *Linear-2* boundary, followed by the *Linear-1* boundary, with lowest separability for the *Nonlinear* boundary (Figure 1D). A similar pattern was found when computing separability using features from a self-supervised deep neural network model (SimCLR; T. Chen et al., 2020; see *Methods*), suggesting that these relationships held even when considering a broader set of image features. The low separability of the *Nonlinear* categories relative to the *Linear-1* and *Linear-2* categories is consistent with the *Nonlinear* boundary being nonlinear in shape space.

Across participants, behavioral accuracy (Figure 1E) was highest for the *Linear-2* task (0.86 ± 0.02; mean ± SEM across 10 participants), followed by the *Linear-1* task (0.83 ± 0.01) and the *Nonlinear* task (0.80 ± 0.01). A repeated measures ANOVA revealed a main effect of task (F_(2,18)_ = 13.22, p < 0.001; p-values obtained using permutation test; see *Methods*), and post-hoc tests showed that accuracy was significantly higher for both of the linear tasks versus the *Nonlinear* task (*Linear-1* vs. *Nonlinear*: t_(9)_ = 2.19, p = 0.024; *Linear-2* vs. *Nonlinear*: t_(9)_ = 4.98, p = 0.002; paired t-tests with permutation; see *Methods*), and higher for the *Linear-2* task versus the *Linear-1 task* (*Linear-1* vs. *Linear-2*: t_(9)_ = −3.00, p = 0.001). This advantage for the *Linear-2* task is consistent with the high relative separability across the *Linear-2* boundary based on image features shown in the previous analysis (Figure 1D). In terms of response times (RTs), a significant main effect of task was also found (F_(2,18)_ = 3.94, p = 0.036; p-values obtained using permutation test). No difference in RTs between the *Linear-1* and *Linear-2* tasks was observed, but RTs were significantly slower for the *Nonlinear* task than the *Linear-1* task (t_(9)_ = −3.08, p = 0.012). In addition to these differences across tasks, we also observed a consistent difference between performance on the easy and hard trials within each task (Figure 1F), which was expected based on the task design. Accuracy was significantly higher on easy versus hard trials within each task (*Linear 1:* t_(9)_ = 11.05, p = 0.002; *Linear-2:* t_(9)_ = 7.88, p = 0.002; *Nonlinear:* t_(9)_ = 15.37, p = 0.002), and RT was significantly faster on easy versus hard trials within each task (*Linear 1:* t_(9)_ = −7.48, p = 0.002; *Linear-2:* t_(9)_ = −9.38, p = 0.002; *Nonlinear:* t_(9)_ = −4.92, p = 0.003).

Next, we examined the neural representations of shape stimuli in each task, under the hypothesis that shape representations would differ across task conditions in accordance with the changing decision boundary. To achieve this we used multivariate classification to analyze single-trial voxel activation patterns from retinotopically defined ROIs (Figure 2). First, we trained a series of binary classifiers to predict the category of the shape shown on each trial, according to each of the three decision boundaries, using data from each task separately (Figure 2A-C). These binary classifiers provide an estimate of the discriminability of shape representations in visual cortex across each of the three decision boundaries, within each task context. Overall, we observed that binary classifier accuracy was highest in early visual areas V1 and V2, and lower in higher visual areas such as LO2 and IPS, although participant-averaged classification accuracy was significantly above chance for every ROI in every task (significance evaluated using a permutation test; FDR corrected; all q < 0.01; see *Methods*). We also observed that accuracy was highest for the *Linear-2* binary classifier (V2 accuracy averaged across tasks: 0.86 ± 0.02; mean ± SEM across 10 participants), followed closely by the *Linear-1* classifier (V2 accuracy averaged across tasks: 0.80 ± 0.02), with lowest accuracy for the *Nonlinear* classifier (V2 accuracy averaged across tasks: 0.72 ± 0.02). However, the overall accuracy of these binary classifiers did not differ significantly across tasks: for each classifier, we performed a two-way repeated measures ANOVA on the classifier values with factors of ROI and task, and found significant main effects of ROI, but no main effects related to task (see Supplementary Table 2 for test statistics).

**Figure 2.**
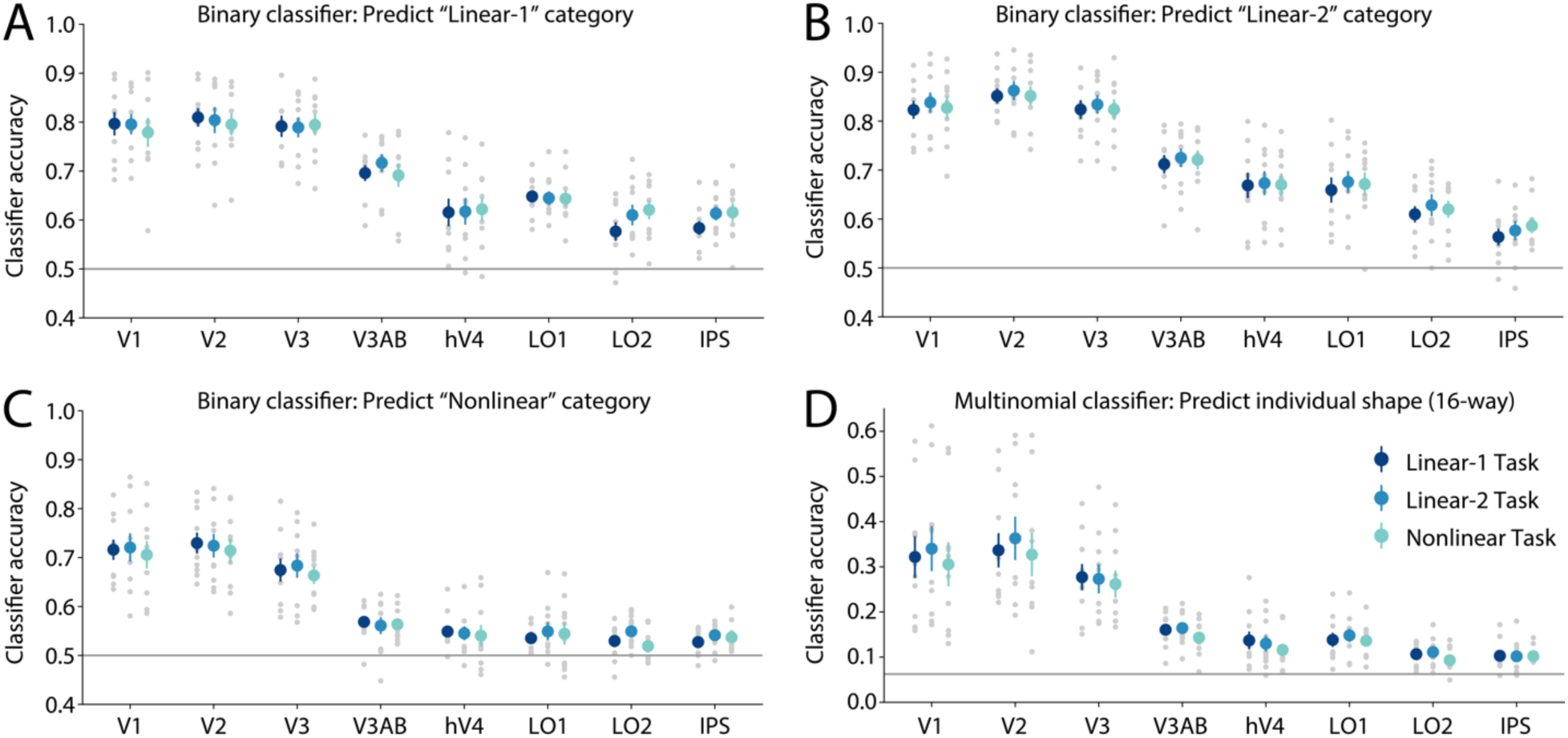
Overall classification accuracy for binary and multinomial classifiers. **(A-C)** A binary logistic regression classifier was trained to predict the category of the shape shown on each trial, according to either the *Linear-1*, *Linear-2*, or *Nonlinear* decision rule. **(D)** A multinomial (16-way) logistic regression classifier was trained to predict the individual shape shown on each trial. In **(A-D)**, classifiers were trained and tested within each task condition separately, training using data from the main grid trials only (i.e. black dots in Figure 1B). Different colors indicate data from different tasks. Plotted values reflect overall prediction accuracy of classifiers for each task and each ROI, computed using trials from the main grid only. Gray dots represent individual participants, colored circles and error bars represent the mean ± SEM across 10 participants, horizontal line indicates chance decoding accuracy (1/2 for binary classifier, 1/16 for multinomial). All classification accuracy values were above chance at the participant-averaged level (FDR corrected, q < 0.01); see *Methods* for more details.

Given that there was no difference in overall binary classifier accuracy across tasks, we next performed a more targeted analysis, based on the hypothesis that task-related differences in category discriminability might be limited to a subset of trials, and therefore would not be measurable when averaging across all trials. Specifically, we predicted stronger effects for shapes nearer to the category boundary versus shapes further from the boundary. To test this, we used the same series of binary classifiers from the previous analysis, but we separated test trials into two groups based on distance to the boundary: “near” trials consisted of the 8 main grid shapes that were closest to the classifier boundary, while “far” trials consisted of the 8 shapes further from the boundary (Figure 3, see diagrams on right side). Note that the “near” group does not include the set of trials that are outside the main grid and closest to the active boundary in each task (i.e., “hard” trials; light gray dots in Figure 1B), but see Figure 8 for discussion of this trial group. We then computed accuracy within each of these trial subsets, using data from the *Linear-1* and *Linear-2* tasks only.

**Figure 3.**
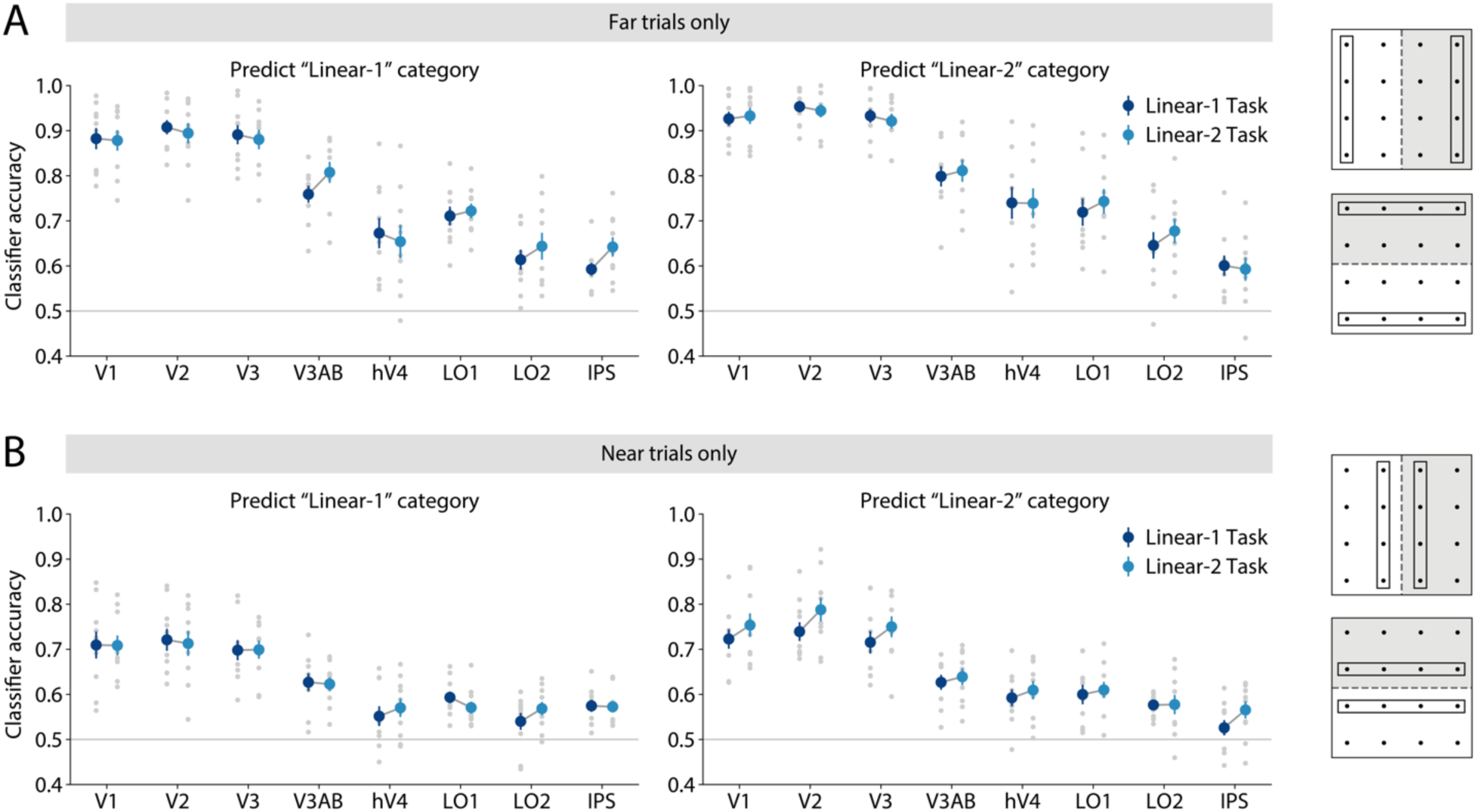
Category separability differs across tasks, only for trials near the decision boundary. Using the binary classifiers that were trained to predict category according to either the *Linear-1* or *Linear-2* decision rule (see Figure 2A-B), we separately computed accuracy using test set trials that were either far or near from the classifier boundary. Each panel shows the results for a different binary classifier (trained to predict either the *Linear-1* or *Linear-2* category), and different colors indicate data from different tasks. **(A)** Accuracy for “far” trials, consisting of the 8 main grid shapes furthest from the classifier boundary (see diagrams on right side of panel for illustration). **(B)** Accuracy for “near” trials, consisting of the 8 main grid shapes nearest to the classifier boundary. In (**A**-**B)**, the gray dots represent individual participants, colored circles and error bars represent the mean ± SEM across 10 participants.

As predicted, this analysis revealed a difference between near and far trials. Classifier accuracy was overall higher for far trials versus near trials, which was expected based on the difference in stimulus discriminability on these trial types. Importantly, we also observed that for near trials only, there was an interaction between classifier boundary and task, such that the accuracy of each classifier appeared higher when the classifier matched the boundary that was currently active in the task. This effect was most pronounced in early areas such as V2. We examined this pattern by performing a three-way repeated measures ANOVA on the classifier accuracy values for near trials, which revealed significant main effects of ROI, Task, and Boundary, as well as a Task x Boundary interaction (ROI: F_(7,63)_ = 65.53, p < 0.001; Task: F_(1,9)_ = 5.37, p = 0.044; Boundary: F_(1,9)_ = 9.33, p = 0.014; Task x Boundary: F_(1,9)_ = 8.99, p = 0.011; p-values obtained using permutation test; see Supplementary Table 3 for complete set of test statistics). We then examined each classifier boundary separately, which showed that across all ROIs, the accuracy of the *Linear-2* classifier for near trials was higher when using data from the *Linear-2* task versus the *Linear-1* task (two-way repeated measures ANOVA; ROI: F_(7,63)_ = 50.00, p < 0.001; Task: F_(1,9)_ = 10.30, p = 0.011; ROI x Task: F_(7,63)_ = 0.83, p = 0.570). At the single ROI level, this difference was significant in V2 (t_(9)_ = −3.27, p = 0.009; paired t-test with permutation; see *Methods*), and V3 (t_(9)_ = −2.80, p = 0.024). However, when examining the accuracy of the *Linear-1* classifier across tasks, no significant difference was observed (two-way repeated measures ANOVA; ROI: F_(7,63)_ = 42.38, p < 0.001; Task: F_(1,9)_ = 0.05, p = 0.828; ROI x Task: F_(7,63)_ = 0.75, p = 0.627). Overall, these results support the idea that on near trials, shape representations may be modified adaptively to become more separable across the task-relevant boundary, particularly during the *Linear-2* task. Notably, performing the same test on the classifier accuracy values from far trials showed no significant interaction between task and classifier boundary (see Supplementary Table 3), suggesting that the modulatory effect of task on visual representations was limited to trials closer to the decision boundary.

To evaluate whether a similar interaction between task, boundary and distance was present for the *Nonlinear* boundary, we performed a similar analysis for the *Nonlinear* binary classifier (Figure 4). Specifically, we computed *Nonlinear* classifier accuracy, separately for trials that were near versus far from the *Nonlinear* decision boundary. In this case, however, we did not observe any consistent differences in classifier accuracy across tasks, for either near trials (two-way repeated measures ANOVA; ROI: F_(7,63)_ = 45.99, p < 0.001; Task: F_(2,18)_ = 0.19, p = 0.829; ROI x Task: F_(14,126)_ = 0.77, p = 0.696), or far trials (two-way repeated measures ANOVA; ROI: F_(7,63)_ = 59.44, p < 0.001; Task: F_(2,18)_ = 1.01, p = 0.380; ROI x Task: F_(14,126)_ = 0.66, p = 0.804).

**Figure 4.**
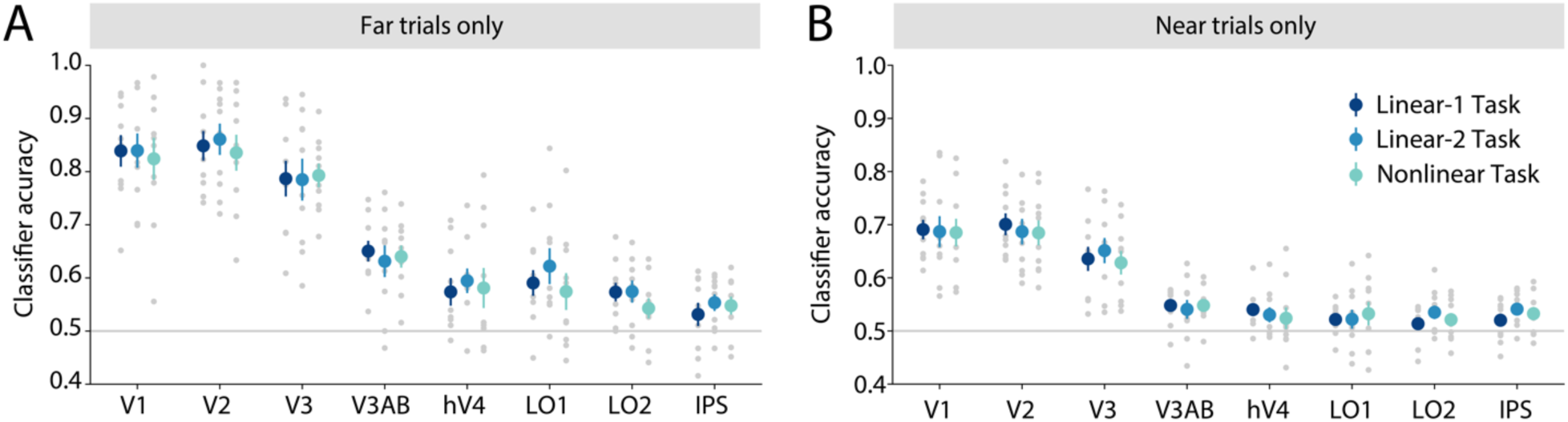
Separability of representations across the *Nonlinear* boundary does not differ significantly across tasks. We computed classifier accuracy for the *Nonlinear* classifier (Figure 2C), separately for trials near versus far from the category boundary. **(A)** Accuracy computed using “far” trials, meaning the four points in the main grid that fell furthest from the two category boundaries (i.e., four corners of the shape space grid). **(B)** Accuracy computed using “near” trials, meaning the 12 points in the main grid that fell nearest to either of the two category boundaries. In (**A**-**B)**, the gray dots represent individual participants, colored circles and error bars represent the mean ± SEM across 10 participants.

Next, we investigated visual cortex representations at a finer level of granularity, by training a 16-way multinomial classifier (Figure 2D). In contrast to the binary classifier analysis, which reduces all stimuli to two discrete categories, this multinomial classifier treats each of the individual shapes as a distinct category, and therefore may be able to pick up on more fine-grained changes to the overall representational space that occur across tasks. As before, we trained and tested this classifier using data from each task separately. We observed that overall 16-way classification accuracy was highest in V2 (16-way accuracy averaged across tasks: 0.34 ± 0.04; mean ± SEM across 10 participants), followed by V1 (0.32 ± 0.05) and V3 (0.27 ± 0.03). Participant-averaged classification accuracy was significantly above chance for every ROI in every task (significance evaluated using a permutation test; FDR corrected; all q < 0.01; see *Methods*).

To characterize the neural shape space, we used the output of the 16-way classifier to compute a confusion matrix for each ROI and for each task, which captures how often the classifier assigned each shape label to each shape in the test dataset (Figure 5; see *Methods*). For V1, this confusion matrix revealed that shape confusability was related to distance in shape space, with the classifier tending to make more errors between shapes that were adjacent in shape space (off-diagonal structure in Figure 5A). This relationship with distance can also be seen by plotting the proportion of predictions as a function of the distance between predicted and actual shape space coordinates (Figure 5B). Importantly, the distances between shape space points were not specified in the construction of the classifier, where all 16 points were treated as independent categories. Thus, the emergence of this structure in the classifier confusion matrix provides evidence for a two-dimensional representation of the shape space grid in V1. A similar pattern was seen in all other ROIs tested.

**Figure 5.**
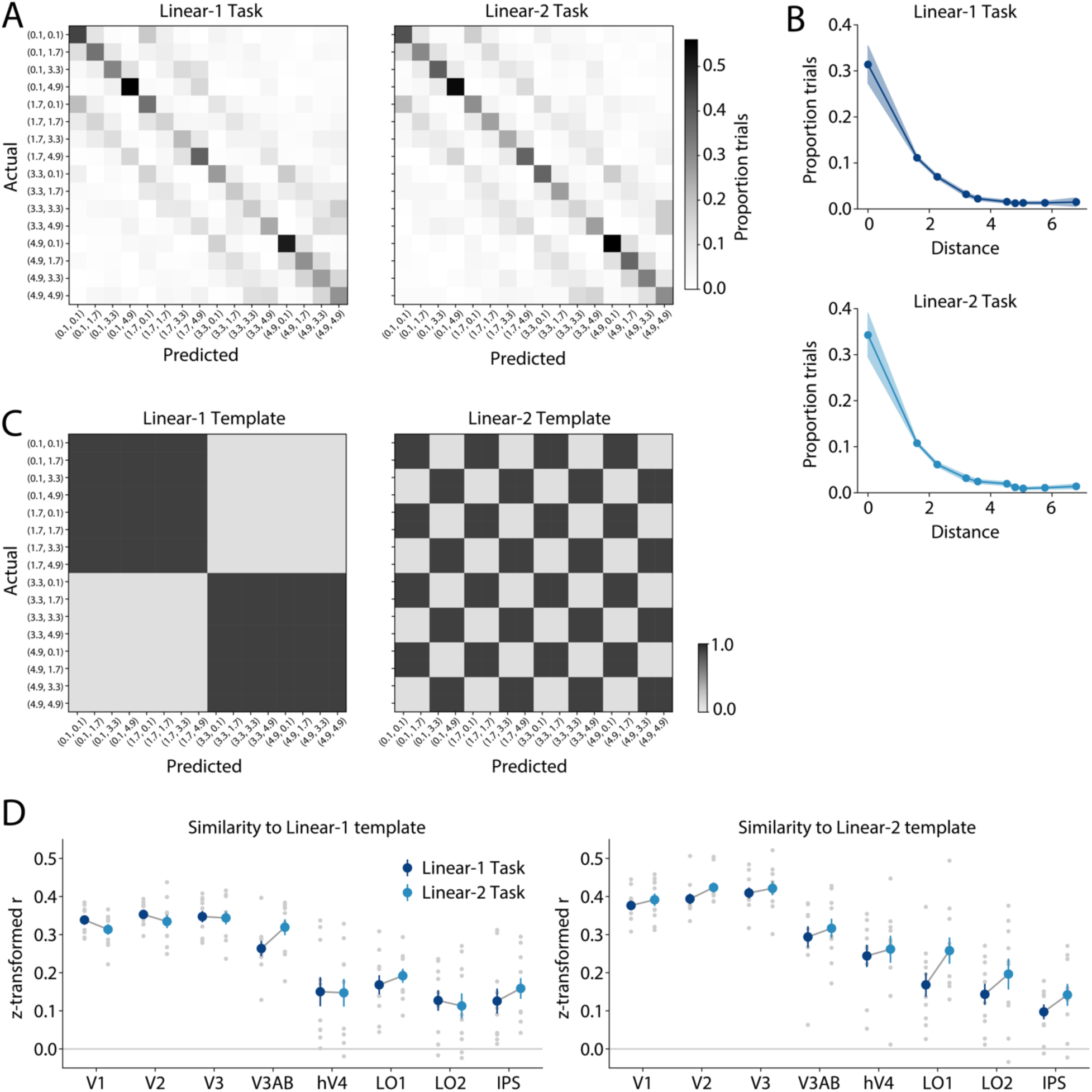
Classifier confusion matrices suggest restructuring of shape representations between the *Linear-1* and *Linear-2* tasks. **(A)** Classifier confusion matrices for V1 in each task, where each row represents the set of trials on which a given shape was actually shown, and the columns represent the proportion of those trials that the classifier predicted as having each of the 16 shape labels (each row sums to 1). Confusion matrices were computed using main grid trials only, and are averaged across 10 participants. **(B)** A simplified view of the classifier confusion data for V1: we computed the proportion of trials on which the actual and predicted shapes were separated by a given distance in shape space. Colored lines and shaded error bars indicate mean ± SEM across 10 participants. **(C)** Template matrices for the *Linear-1* and *Linear-2* tasks, representing the pattern of confusability expected for a perfect binary representation of each decision boundary. In A and C, the axis labels are coordinate pairs which represent the position of stimuli in shape space: (axis 1 coordinate, axis 2 coordinate). These are analogous to the x and y coordinates in Figure 1B. The *Linear-1* template distinguishes stimuli based on their axis 1 coordinate (x), while the *Linear-2* template distinguishes stimuli based on their axis 2 coordinate (y). **(D)** The similarity (Pearson correlation coefficient, z-transformed) between actual and template confusion matrices for each task and each ROI. Gray dots represent individual participants, colored circles and error bars represent the mean ± SEM across 10 participants. See Supplementary Figure 1 for an analogous analysis using a template for the *Nonlinear* task.

Next, we examined how well the neural shape space measured in each task aligned with each decision rule. To examine this, we first constructed “template” confusion matrices for the *Linear-1* and *Linear-2* boundaries, where each template had 1 for shape pairs that were on the same side of the category boundary for that task and 0 for shape pairs that were on different sides (Figure 5C). We then correlated these template matrices with the real confusion matrices for each task (Figure 5D). This analysis revealed that the similarity of confusion matrices to each template differed depending on task. A three-way repeated measures ANOVA on the z-transformed template similarity values showed main effects of ROI and Template, as well as a significant ROI x Template interaction and a significant Task x Template interaction (ROI: F_(7,63)_ = 46.42, p < 0.001; Task: F_(1,9)_ = 8.06, p = 0.020; Template: F_(1,9)_ = 21.05, p = 0.001; ROI x Task: F_(7,63)_ = 1.41, p = 0.217; ROI x Template: F_(7,63)_ = 3.25, p = 0.004; Task x Template: F_(1,9)_ = 8.89, p = 0.015; ROI x Task x Template: F_(7,63)_ = 0.97, p = 0.461; p-values obtained using permutation test; see *Methods*). Evaluating the similarity values for each template separately, we found that across all ROIs, the *Linear-2* template was significantly more similar to confusion matrices computed from the *Linear-2* task versus the *Linear-1* task (two-way repeated measures ANOVA; ROI: F_(7,63)_ = 31.99, p < 0.001; Task: F_(1,9)_ = 15.62, p = 0.003; ROI x Task: F_(7,63)_ = 0.97, p = 0.467). Post-hoc tests showed that the difference in similarity to the *Linear-2* template between the *Linear-2* and *Linear-1* tasks was significant in LO1 (t_(9)_ = −2.93, p = 0.007; paired t-test with permutation; see *Methods*). These findings suggest that shape representations in LO1 were more aligned with the *Linear-2* template when the *Linear-2* boundary was relevant than when it was irrelevant for the present task. However, the similarity of confusion matrices to the *Linear-1* template did not differ significantly across tasks (two-way repeated measures ANOVA; ROI: F_(7,63)_ = 32.57, p < 0.001; Task: F_(1,9)_ = 0.49, p = 0.502; ROI x Task: F_(7,63)_ = 1.53, p = 0.175). Additionally, when we constructed a template for the *Nonlinear* task, we did not observe a difference in the similarity of confusion matrices to the *Nonlinear* template across tasks (Supplementary Figure 1). Together, these results suggest that shape representations in visual cortex during our task may reorganize in a way that reflects the current decision boundary and shifting cognitive demands.

As in the binary classifier analysis, we then asked whether these representational changes were more pronounced for shapes nearer to the category boundary than shapes further from the boundary. We again divided the trials into near and far groups based on distance to the boundary. To measure the category separability of shapes in each of these distance bins, we computed a continuous measure we refer to as classifier confidence (Figure 6). Confidence is a single-trial measure, computed with respect to each of the category boundaries separately, and was computed by taking the output of the 16-way classifier described above and comparing the total probability assigned by the classifier to points on each side of each boundary. Larger positive values indicate higher separability of shapes across the boundary of interest. We refer to these measures, with respect to each boundary, as *Linear-1* confidence, *Linear-2* confidence, and *Nonlinear* confidence.

**Figure 6.**
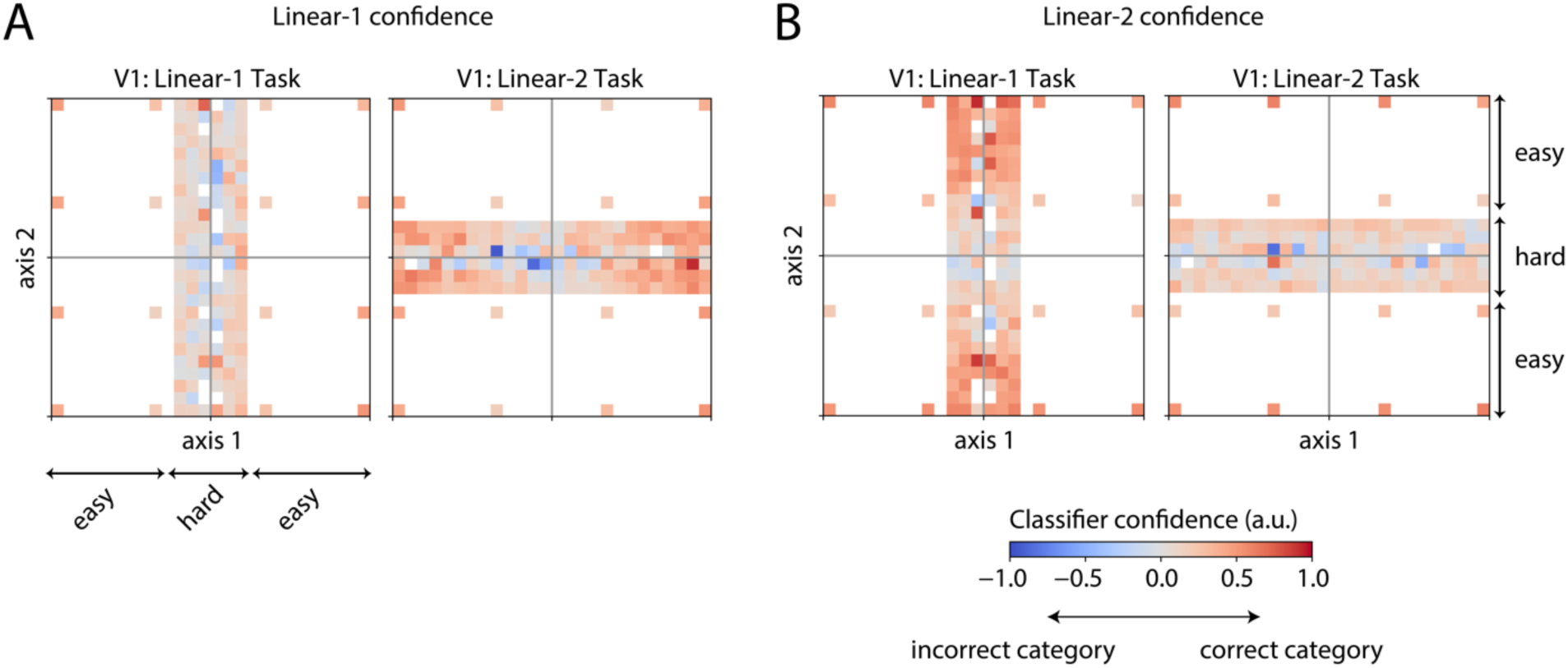
Illustration of how classifier “confidence” was computed with respect to each binary decision boundary. **(A)** *Linear-1* confidence, or confidence with respect to the *Linear-1* category boundary, was computed based on the difference between the total probability assigned by the 16-way classifier to each side of the boundary (see *Methods*). Left and right panels represent data from V1 in the *Linear-1* and *Linear-2* tasks, respectively, averaged across all participants. In each of the plots, each square represents a bin of shape space positions in the test dataset, and the color indicates the average confidence assigned to the correct category for that test trial (red) versus the incorrect category (blue). Arrows labeled “easy” and “hard” indicate the trial types, as in Figure 1B; the “hard” trial group was only used to generate Figure 8. **(B)** Same as A, but showing *Linear-2* confidence. An analogous procedure was also used to compute *Nonlinear* confidence; see *Methods*.

We then compared *Linear-1* confidence and *Linear-2* confidence across the *Linear-1* and *Linear-2* tasks (Figure 7). Overall, both types of confidence were highest for trials furthest from the boundary (Figure 7A), followed by near trials (Figure 7B). This pattern is expected given that shapes further from the boundary are more distinctive from one another, while shapes nearer to the boundary are more ambiguous. In addition, this analysis revealed effects of task condition that differed for near and far trials. For trials in the far group, a three-way repeated measures ANOVA showed main effects of ROI and confidence boundary (i.e., *Linear-1* confidence versus *Linear-2* confidence), but no main effect of task or interaction between task and boundary (Supplementary Table 4), suggesting that discriminability of shapes across the *Linear-1* and *Linear-2* boundaries did not differ across tasks for this group of trials. For the near trials, however, there was also a significant interaction between task and boundary (Supplementary Table 4). When each boundary was examined separately for each of these trial groups, we found a main effect of task on *Linear-2* confidence for the near trials (two-way repeated measures ANOVA on near trials; ROI: F_(7,63)_ = 30.05, p < 0.001; Task: F_(1,9)_ = 13.65, p = 0.005; ROI x Task: F_(7,63)_ = 0.36, p = 0.925), with *Linear-2* confidence showing higher values for the *Linear-2* task, across all ROIs, than the *Linear-1* task. As with the previous analyses, the effect of task was larger for the *Linear-2* boundary than for the *Linear-1* boundary – there was no main effect of task seen for the *Linear-1* confidence values for near trials (ROI: F_(7,63)_ = 23.58, p < 0.001; Task: F_(1,9)_ = 0.10, p = 0.757; ROI x Task: F_(7,63)_ = 0.62, p = 0.751). As a further test, we also performed a version of this classifier confidence analysis using the output of the simpler binary classifiers presented earlier (Supplementary Figure 2). This revealed the same pattern of results, namely an interaction between the classifier boundary and the task, in which *Linear-2* confidence values were significantly higher when computed from the *Linear-2* task versus the *Linear-1* task. This indicates that the difference in classifier confidence across tasks is not dependent on the classifier training method used.

**Figure 7.**
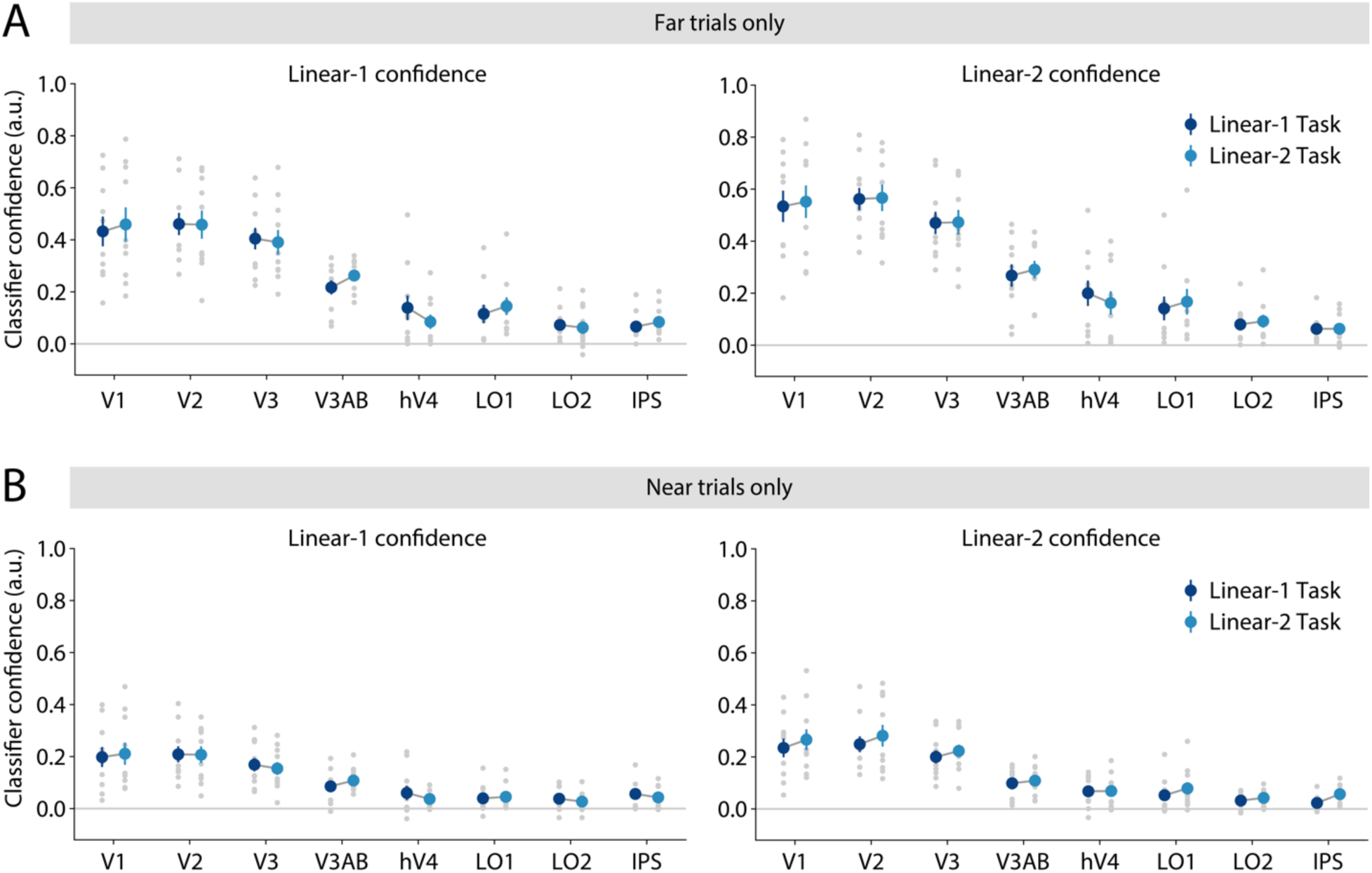
Discriminability of *Linear-1* and *Linear-2* shape categories depends on task and proximity to category boundaries. To obtain a continuous estimate of shape category discriminability, we used our 16-way multinomial classifier (see Figure 2D) to compute classifier confidence toward the correct binary category on each trial (see Figure 6). Confidence was computed with respect to the *Linear-1* categorization boundary (*Linear-1* confidence; left) or the *Linear-2* categorization boundary (*Linear-2* confidence; right). **(A)** Confidence computed using “far” trials, meaning the 8 points in the main grid that fell furthest from the category boundary of interest. **(B)** Confidence computed using “near” trials, meaning the 8 points in the main grid that fell nearest to the boundary of interest. In (**A**-**B)**, the gray dots represent individual participants, colored circles and error bars represent the mean ± SEM across 10 participants. For an analogous version of this analysis based on a binary classifier, see Supplementary Figure 2.

In addition to comparing confidence across the two linear boundaries, we measured *Nonlinear* confidence for the far and near trials in each task (Supplementary Figure 3). As before, confidence values tracked the distance of shapes from the boundary, with highest overall confidence observed for far trials. In contrast to the results with *Linear-2* confidence, however, *Nonlinear* confidence did not show any significant differences across tasks.

Finally, we evaluated whether the discriminability of shape representations across the relevant category boundary in each task was associated with behavioral performance. To test this, we compared classifier confidence for correct versus incorrect trials: focusing here on only the “hard” trials (see light gray points in Figure 1B), because these had the highest rate of incorrect responses. To ensure a fair comparison across correct and incorrect trials, we used bootstrap resampling to match the distribution of stimulus positions sampled in each group of trials; see *Methods* for details. As shown in Figure 8, this analysis revealed a significant difference in classifier confidence between correct and incorrect trials in both the *Linear-2* and the *Nonlinear* tasks, with confidence tending to be higher for correct trials than incorrect trials, particularly in early areas V1, V2, and V3. A two-way repeated measures ANOVA with factors of ROI and correctness revealed a significant main effect of correctness for both the *Linear-2* and *Nonlinear* tasks, and a significant interaction between ROI x correctness for the *Nonlinear* task (*Linear-2;* ROI: F_(7,63)_ = 10.21, p < 0.001; Correctness: F_(1,9)_ = 6.33, p = 0.031; ROI x Correctness: F_(7,63)_ = 1.81, p = 0.099; *Nonlinear*; ROI: F_(7,63)_ = 7.55, p < 0.001; Correctness: F_(1,9)_ = 8.68, p = 0.016; ROI x Correctness: F_(7,63)_ = 2.82, p = 0.011; p-values obtained using permutation test; see *Methods*). At the individual ROI level, confidence was significantly higher for correct versus incorrect trials in V1 during both the *Linear-2* and the *Nonlinear* tasks (*Linear-2;* t_(9)_ = 3.62, p = 0.007; *Nonlinear;* t_(9)_ = 3.39, p = 0.008; paired t-test with permutation; see *Methods*), and in V2 during the *Linear-2* task (t_(9)_ = 2.91, p = 0.022). The *Linear-1* task showed no significant differences in confidence for correct versus incorrect trials (ROI: F_(7,63)_ = 4.90, p < 0.001; Correctness: F_(1,9)_ = 0.40, p = 0.543; ROI x Correctness: F_(7,63)_ = 0.98, p = 0.453). These results indicate that the separability of shape representations in early visual cortex across the task-relevant category boundary was associated with behavioral performance, at least for two out of three categorization tasks.

**Figure 8.**
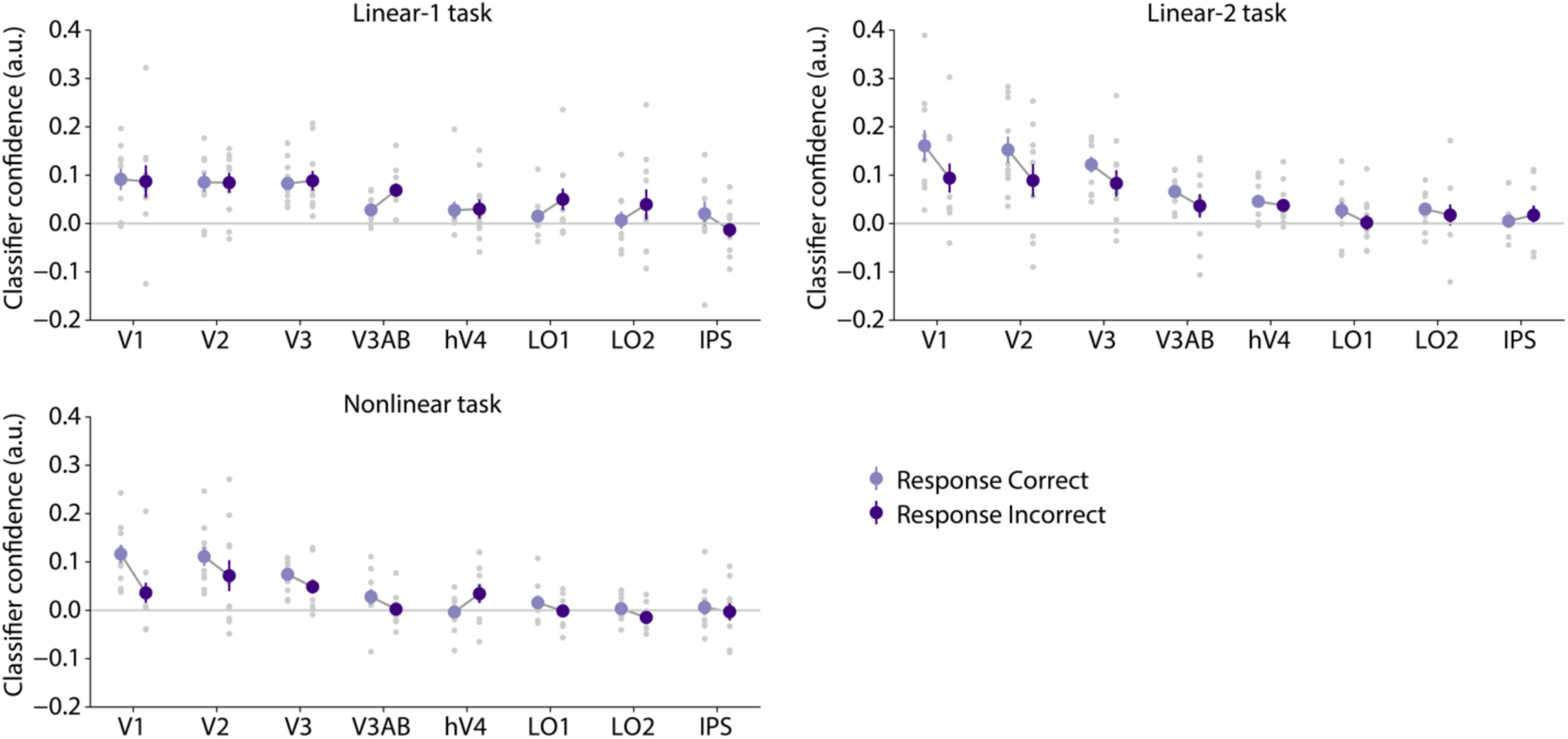
Task-relevant shape categories are more discriminable on correct versus incorrect trials. In each task, classifier confidence was computed with respect to the relevant category boundary for that task. Confidence was computed using “hard” trials only (those not on the main grid, and nearest the relevant boundary), separately for trials with correct and incorrect behavioral responses. The set of shape space positions sampled on correct and incorrect trials was matched using resampling to ensure that the effect was not driven by stimulus differences; see *Methods* for details. Gray dots represent individual participants, colored circles and error bars represent the mean ± SEM across 10 participants.

## Discussion

Our goal was to determine whether and how human visual cortex representations of shape stimuli are adaptively modulated when switching between distinct task contexts. To test this, we trained participants to perform a categorization task on shape silhouette stimuli within a two-dimensional shape space (Figure 1). Participants categorized shapes according to different categorization rules (*Linear-1, Linear-2, Nonlinear*) on interleaved fMRI scanning runs, and we used multivariate decoding to explore how neural representations shift based on decision rules and the relative positions of shapes within the two-dimensional stimulus space. We showed that the discriminability of shapes across each linear boundary, as measured by classifier accuracy and classifier confidence, was higher when that boundary was relevant to the current task.

These effects were most pronounced in early areas V1-V3, and were strongest for shapes located nearest to the active categorization boundary (Figure 3, Figure 7). We also used a confusion matrix analysis to show that shape representations became more aligned with the *Linear-2* boundary when participants were performing the *Linear-2* task versus the *Linear-1* task, with the largest effect observed in LO1 (Figure 5). Finally, we showed that the discriminability of shapes across relevant category boundaries was higher on correct versus incorrect trials, indicating a link with behavioral task performance (Figure 8). Together, these results demonstrate that performance of a categorization task with a dynamically changing task boundary is accompanied by changes to neural representations in human visual cortex.

The average accuracy of our classifiers, across tasks, was highest in V2 followed by V1 and V3. This high decoding accuracy in early areas is surprising in light of earlier work suggesting that higher visual areas like ITC and LOC encode shapes similar to ours (i.e., radial frequency components (RFC)-defined silhouettes) in a way that matches perceptual similarity (Drucker & Aguirre, 2009; Op de Beeck et al., 2001), and that LOC is critically involved in shape computations (Vinberg & Grill-Spector, 2008). Work in non-human primates also indicates that neurons in ITC, as well as in V4, are more strongly tuned for shape and contour than neurons in V1 (Connor et al., 2007; DiCarlo & Maunsell, 2000; Pasupathy & Connor, 1999; Tanaka, 1993, 1996). One reason for our observation of higher decoding accuracy in early areas is that our stimuli were silhouettes presented at a fixed size and position, so invariance to size or position was not required to encode them accurately. As a result, fine-grained retinotopic and orientation tuning in areas like V1-V3 was likely sufficient to encode the shapes with high accuracy, without the need for an explicit – or invariant – contour or shape representation. Importantly, the goal of our experiment was not to measure abstract representations of shape or contour *per se* but to measure how visual representations change in accordance with dynamically varying decision boundaries, and our relatively simple stimulus set was appropriate for this goal.

The effects of task context on classifier accuracy and classifier confidence (Figure 3, Figure 7), as well as association of classifier confidence with behavioral performance (Figure 8), also tended to be strongest in early visual areas. This advantage for early areas may be due in part to the higher signal-to-noise ratio (SNR) of decoding accuracy in V1-V3, but it may also suggest that representations in these areas are particularly important for performance of our decision task. The findings of strong task-dependent effects in early retinotopic areas align with recent rodent studies, which show that representations within sensory areas contain information pertinent to task goals, motor outcomes, and prior knowledge about sensory environments (Ebrahimi et al., 2022; Findling et al., 2023; Mimica et al., 2023; Niell & Stryker, 2010; Stringer et al., 2019). Extending these findings, our study demonstrates that human visual areas are more actively involved with decision-related computation than previously thought. Our results demonstrate that human sensory areas not only code for temporally varying task contexts but also dynamically integrate this information with incoming sensory inputs to optimize decision processes. This observation challenges the traditional view that sensory areas are primarily dedicated to basic sensory processing, suggesting a more multifaceted role in cognitive computation.

A plausible mechanism for guiding dynamic task coding and context-dependent representation of sensory inputs in humans may involve the deployment of selective attention. By flexibly prioritizing processing of relevant stimulus features based on current task goals, attention may guide the integration of sensory information with shifting task demands. Specifically, our observed task-dependent effects in early retinotopic areas are consistent with the literature on feature-based attention, which has shown that directing attention to simple visual features can modulate representations in early visual cortex (X. Chen et al., 2012; Foster & Ling, 2022; Gundlach et al., 2023; Jehee et al., 2011; Liu et al., 2003, 2007; Martinez-Trujillo & Treue, 2004; Mirabella et al., 2007; Saenz & Boynton, 2003; Serences & Boynton, 2007; Treue & Maunsell, 1996, 1999; Yoo et al., 2022). By modulating neurons coding for perceptual features that differentiate between categories, feature-based attention could provide a mechanism for improving the separability of different stimulus categories (Navalpakkam & Itti, 2007; Scolari et al., 2012; Scolari & Serences, 2009). Our result of early modulations is also consistent with Ester et al. (2020), who found biases in orientation representations that were related to categorization, although their paradigm used a single category boundary as opposed to a dynamically updated boundary.

Importantly, however, our experiment differs from typical paradigms for studying feature-based attention (Martinez-Trujillo & Treue, 2004; Saenz & Boynton, 2003; Treue & Maunsell, 1996; Treue & Maunsell, 1999; Desimone & Duncan, 1995) in that participants were not cued explicitly to a single elementary feature dimension (such as orientation or motion direction), and instead were required to categorize stimuli along axes in an abstract shape space. Within the shape space, simple features like a single orientation or retinotopic position are not sufficient to determine the category of a shape, so information must be integrated over multiple areas of the image and multiple low-level feature dimensions in order to solve the task. In this light, one hypothesis for our observed results is that during each task, a subset of the neurons within early visual cortex are tuned for feature combinations that are diagnostic of the relevant category distinction. These subpopulations may be tuned for specific retinotopic regions of the image, features like orientation or curvature, or combinations of these properties. Top-down modulations may then selectively target these particular subpopulations, leading to an increase in shape discriminability at the population level. In this respect, our results go beyond existing knowledge on selective attention, by showing that a mechanism similar to feature-based attention, perhaps combined with spatial attention, may operate in visual cortex within the context of a more complex, abstract decision-making task.

Relatedly, other work using more complex stimuli such as three dimensional objects and human bodies has also shown feature-based attention effects in higher visual areas such as LOC and the extrastriate body area (EBA), as opposed to early visual cortex (Jackson et al., 2017; Thorat & Peelen, 2022). As discussed earlier, the fact that we saw larger effects in early visual areas versus higher areas may be due to the fact that our task did not require position-invariant representations of shape or contour. Interestingly, Jackson et al. (2017) also examined early visual areas in their study of three-dimensional object coding, and found that while LOC encoded more information about a task-relevant object dimension, no such effect was found in early visual areas. One possible explanation for this is that our stimuli subtended a large portion of the visual field, with the most category-diagnostic features distributed across a range of retinotopic positions, while in the stimuli used by Jackson et al., the task-relevant stimulus features were localized to a small region of the image. This difference in spatial distribution, and possibly the allocation of spatial attention, may explain why we observed task-related modulations in early retinotopic cortex while Jackson et al. did not. More generally, these observations may indicate that attentional modulations in V1-V3 are most important for task performance when stimuli are relatively simple and require fine-grained spatial detail (e.g., oriented gratings, two-dimensional silhouettes in our task), than when stimuli are more complex and require position invariance. In keeping with this idea of attention adapting dynamically to the most informative features for a task, a recent behavioral study demonstrated that feature-based attention is adaptively allocated according to experience with the variance of feature distributions (Witkowski & Geng, 2022). Our findings extend these prior studies by demonstrating feature-based attention as a potential mechanism for effectively integrating sensory information with changing task requirements within human sensory cortex.

Despite the relatively low classifier accuracy values that were observed in higher areas, we did observe a significant effect of task-relevance in LO1 based on the confusion matrix analysis in Figure 5. In this analysis, we demonstrated that classifier confusion matrices from LO1 were more aligned with the *Linear-2* task template during the *Linear-2* task versus the *Linear-1* task. The divergence of this finding from our classifier accuracy and confidence analyses, in which early areas showed larger task effects than LO1, may indicate that the nature of representational changes in LO1 across categorization tasks differs from the changes in V1-V3. Specifically, the confusion matrix analysis tests the hypothesis that shape representations in each task become more aligned with a binary, categorical code, and tests this hypothesis using all trials together. The classifier accuracy and confidence analyses, on the other hand, test for an increase in category discriminability specifically for trials that are near the boundary. In this light, one interpretation is that context-related changes in early areas reflect subtle changes in discriminability that are limited to the area near the category boundary. These subtle changes allow the overall structure of the representational space to be largely maintained across tasks in a stable sensory code. On the other hand, changes in LO1 may reflect a more dramatic restructuring of sensory codes into a format that resembles a binary or categorical code for each task. Such a difference would be consistent with LO1 being a higher visual area more closely aligned with decision processes than early areas. In addition to this, the confusion matrix analysis captures changes to the relationship between all 16 shapes in the main shape space grid, including pairs on the same side of the boundary, while the classifier accuracy and confidence analyses only capture the discriminability of shapes across the category boundary. Based on this, another (non-exclusive) hypothesis is that the changes in LO1 from the *Linear-1* task to the *Linear-2* task are primarily driven by re-structuring of shape representations within a given category (i.e. “acquired equivalence”; Goldstone, 1994) as opposed to an increase in discriminability across the boundary. Further experiments will be needed to evaluate these possibilities.

When classifier accuracy and confidence values were broken down based on proximity to the category boundary, we observed the largest effects of categorization task on confidence for stimuli nearest the boundary, and no effect of task for the furthest stimulus positions. This scaling of categorization effects with proximity to the boundary is consistent with a previous fMRI experiment (Ester et al., 2020) as well as past behavioral experiments (Ashby & Maddox, 2005; Goldstone, 1994, 1998; Livingston et al., 1998; Newell & Bülthoff, 2002). These convergent findings suggest that top-down modulatory effects in early visual cortex are strengthened on trials with higher category ambiguity, facilitating perceptual discrimination of these challenging stimuli. Importantly, our results also build on these past findings by demonstrating an increase in the discriminability of representations near the decision boundary during a task that requires flexible switching between multiple decision boundaries.

Task context had more consistent effects on discriminability with respect to the *Linear* tasks compared to the *Nonlinear* task, with no significant difference across tasks observed for *Nonlinear* classifier accuracy (Figure 4). This difference may be due to the fact that the *Nonlinear* task required using a non-linear decision boundary. The non-linear boundary was more challenging behaviorally, as demonstrated by the slower RTs and lower accuracy observed in the *Nonlinear* task compared to the *Linear-1* and *Linear-2* tasks, which is also consistent with a past report showing that a quadrant task with similar stimuli was more challenging for macaques to learn than a linear rule (Op de Beeck et al., 2001). Notably, our image similarity analysis (Figure 1D) suggested an even more dramatic difference in difficulty between the *Nonlinear* task and the *Linear* tasks, compared to the modest difference seen behaviorally. This may suggest that human observers used a more complex strategy to solve the *Nonlinear* task, allowing them to do relatively well on the *Nonlinear* task despite the low separability of the *Nonlinear* categories in image space. For example, they might have first identified the quadrant each shape belonged to, then mapped this quadrant to a category label using an abstract rule.

In terms of our classifier results, the non-linearity of the boundary may also explain the lack of a consistent task-related modulation of *Nonlinear* discriminability in visual cortex. It is possible that while top-down mechanisms are capable of selectively enhancing representations along one continuous axis in a perceptual space, such a mechanism does not exist for non-linear boundaries. Interestingly, although we did not observe a task-related modulation of *Nonlinear* confidence, we observed a significant within-task association of *Nonlinear* confidence with behavioral performance (Figure 8). One explanation for this difference is that a different set of trials is used for each analysis – the association of confidence with behavioral performance was computed using hard trials only, while the task-related effect was assessed using easy trials only. We did not examine task-related effects on classifier confidence for hard trials here, due to the fact that hard trials sampled different portions of the stimulus space in each task (this was an intended property of the experimental design; see Figure 1B), which made it challenging to obtain fair, stable comparisons of confidence across tasks for these trials. However, it is possible that if sufficient trials had been collected for positions closer to the *Nonlinear* boundary in each task, a task-related enhancement of *Nonlinear* category coding may have been measurable. At the same time, the difference in outcomes between these analyses may also indicate that while discriminability of shapes across the *Nonlinear* boundary does not differ across task contexts, there is variability in the quality of representations across trials within the *Nonlinear* task, and this variability is associated with behavioral performance.

Comparing the two *Linear* tasks, we observed higher SNR for discriminating stimuli across the *Linear-2* boundary than the *Linear-1* boundary (i.e., higher average accuracy of binary classifier across the *Linear-2* boundary, and higher values of similarity to *Linear-2* template, across all tasks). We also observed more consistent effects of task relevance on *Linear-2* accuracy, template similarity, and confidence than the analogous measures with respect to *Linear-1.* Finally, we did not observe any association of *Linear-1* confidence with behavioral performance, though such an effect was observed for *Linear-2* and *Nonlinear* confidence. These findings may be related to the difference in perceptual separability, as measured by our image similarity analyses, between the *Linear-1* and *Linear-2* categories (Figure 1D). The *Linear-2* boundary, across which shapes are more perceptually distinctive, may also be a more effective target of context-dependent processing via selective attention mechanisms. At the same time, however, we note that several of our analyses also revealed a significant interaction between task and classifier boundary (Figure 3B, Figure 5D, Figure 7B), which indicates that there is not simply an increase in signal-to-noise ratio from the *Linear-1* to *Linear-2* task that drives the observed effects, but a specific, task-dependent enhancement of *Linear-2* category separability during the *Linear-2* task. Taken together, these findings may indicate an asymmetry in the allocation of attention to different dimensions within our shape space, in a way that reflects physical properties of the stimuli.

Overall, our findings provide evidence for context-dependent modulations of neural representations in early visual cortex, and show that these effects differ in accordance with temporally shifting task demands. Shape representations were modified to support discrimination of currently-relevant shape categories, with effects that were strongest for stimuli near the decision boundary. Moreover, these effects were associated with task performance. These results may indicate that visual cortex plays an active computational role in the flexible categorization of stimuli, providing new insight into how we organize knowledge about visual stimuli in the face of changing behavioral requirements.

## Materials & Methods

### Human participants

Ten (10) participants were recruited from the UCSD community, and were adults having normal or corrected-to-normal vision. Participants were between the ages of 24 and 33 (mean = 28.2, std = 3.0), and 7 out of 10 were female. The protocol for this study was approved by the Institutional Review Board at UCSD, and all participants provided written informed consent. As part of this experiment, each participant took part in one behavioral training session lasting approximately 1 hour, for which they were compensated at a rate of $10/hour and three scanning sessions each lasting approximately 2 hours, for which they were compensated at a rate of $20/hour. During each scanning session for this experiment, participants also performed several runs of a n-back (repeat detection) task on the same stimuli used in our main task (see *Main task design*). Data from this task are not analyzed here but are included in our full open dataset (see *Data availability*). Each participant also participated in a separate retinotopic mapping scan session; for eight participants this retinotopic mapping session was performed as part of an earlier experiment and for the remaining two it was performed just prior to the start of the present experiment.

### Acquisition of MRI data

All magnetic resonance imaging (MRI) scanning was performed at the UC San Diego Keck Center for Functional Magnetic Resonance Imaging. For the first 7 participants, we used a General Electric (GE) Discovery MR750 3.0T scanner, and for the latter 3 participants, we used a Siemens MAGNETOM Prisma 3.0T scanner. Given that all manipulations were within-subject, we combined data across scanners.

We first discuss the protocols that were used for the GE scans: We used a Nova Medical 32-channel head coil (NMSC075-32-3GE-MR750) to acquire all functional echo-planar imaging (EPI) data, using the Stanford Simultaneous Multislice (SMS) EPI sequence (MUX EPI), with a multiband factor of 8 and 9 axial slices per band (total slices = 72; 2 mm^3^ isotropic; 0 mm gap; matrix = 104 x 104; field of view = 20.8 cm; repetition time/time to echo [TR/TE] = 800/35 ms; flip angle = 52°; inplane acceleration = 1). To perform image reconstruction and un-aliasing we used reconstruction code from the Stanford Center for Neural Imaging, on servers hosted by Amazon Web Services. The initial 16 TRs collected at sequence onset were used as reference images in order to transform data from k-space to image space.

For the Siemens scans: We used a Siemens 32-channel head coil (Siemens Medical Solutions, Malvern, PA) to acquire all functional EPI data. Functional runs used a multiband acceleration factor of 4 (slices = 68; 2.5 mm^3^ isotropic; 0 mm gap; matrix = 100 x 100; field of view = 25.0 cm; repetition time/time to echo [TR/TE] = 1300/32.60 ms; flip angle = 50°; phase-encoding direction A>>P).

In addition, for both types of scanners, a set of two “topup” datasets (17s each) were collected using forward and reverse phase-encoding directions. For the GE scans, we collected one set of topups at the halfway point of the session, and for the Siemens scans, we collected 2-3 sets of topups that were evenly distributed through the session. These runs were used to correct for distortions in the EPI sequences from the same session using topup functionality (Andersson et al., 2003) in the FMRIB Software Library (FSL; Jenkinson et al., 2012).

In addition to the functional data, we also collected a high-resolution anatomical scan for each participant as part of that participant’s retinotopic mapping session. This anatomical T1 image was used for segmentation, flattening, and delineation of the retinotopic mapping data. For five out of the ten participants, we acquired this anatomical scan using the same 32 channel head coil used for functional scanning, and for the remaining five participants, we used an in vivo eight-channel head coil. Anatomical scans were acquired using accelerated parallel imaging (GE ASSET on a FSPGR T1-weighted sequence; 1 x 1 x 1 mm^3^; 8136 ms TR; 3172 ms TE; 8° flip angle; 172 slices; 1 mm slice gap; 256 x 192 cm matrix size). When the 32-channel head coil was used, anatomical scans were corrected for inhomogeneities in signal intensity using GE’s ‘phased array uniformity enhancement’ (PURE) method.

### Preprocessing of functional MRI data

Preprocessing of functional data was performed using tools from FSL and FreeSurfer (available at http://www.fmrib.ox.ac.uk/ fsl and https://surfer.nmr.mgh.harvard.edu). We first performed cortical surface gray-white matter volumetric segmentation of the anatomical T1 scans for each participant, using the recon-all function in FreeSurfer (Dale et al., 1999). The segmented T1 data were then used to define cortical meshes on which we defined retinotopic ROIs (see next section for details). We also used the anatomical T1 data in order to align multi-session functional data to a common space for each participant. This was performed by using the first volume of the first scan for each session as a template, and using this template to align the entire functional session to the anatomical scan for each participant. We used the manual and automatic boundary-based registration tools in FreeSurfer to perform co-registration between functional and anatomical data (Greve & Fischl, 2009), then used the resulting transformation matrix and FSL FLIRT to transform all functional data into a common space (Jenkinson et al., 2002; Jenkinson & Smith, 2001). Next, we used FSL MCFLIRT to perform motion correction (Jenkinson et al., 2002), with no spatial smoothing, with a final sinc interpolation stage, and 12° of freedom. Finally, we performed de-trending to remove slow drifts in the data using a high-pass filter (1/40 Hz cutoff).

Following these initial preprocessing stages, we z-scored the data within each scan run on a per-voxel basis to correct for differences in mean and variance across runs. This and all subsequent analyses were performed using Python 3.7.10 (Python Software Foundation, Wilmington, DE). Next, we obtained a single estimate for each voxel’s activation on each trial by averaging the time series over a window spanning from 3.2-5.6s (4-7 TRs) following image onset (for subjects S01-S07, who were scanned with a 0.8s TR), or from 2.6-6.5s (2-5 TRs) following image onset (for subjects S08-S10, who were scanned with a 1.3s TR). See *Main task design* for more details on task timing and procedure. We then extracted data from voxels within several regions of interest (ROIs; see next section) that were used for subsequent analyses.

### Retinotopic ROI definitions

We defined several retinotopic visual ROIs: V1, V2, V3, V3AB, hV4, LO1, LO2, and IPS, following previously identified retinotopic mapping procedures (Engel et al., 1997; Jerde & Curtis, 2013; Sereno et al., 1995; Swisher et al., 2007; Wandell et al., 2007; Winawer & Witthoft, 2015, Mackey et al., 2017). We combined all intraparietal sulcus (IPS) subregions (IPS0, IPS1, IPS2, IPS3), into a single combined IPS ROI, as this led to improved classifier accuracy relative to the individual sub-regions. For 8 out of 10 participants (all except S08 and S09), retinotopic mapping stimuli consisted of black-and-white contrast reversing checkerboard stimuli that were configured as a rotating wedge (10 cycles, 36 s/cycle), expanding ring (10 cycles, 32 s/cycle), or bowtie shape (8 cycles, 40 s/cycle). During the rotating wedge task, a contrast detection task (detecting dimming events approximately every 7.5 s) was used to encourage covert attention to the stimulus. Average accuracy on this task was 76.75 ± 4.01% (mean ± SEM across 8 participants). The stimulus had a maximum eccentricity of 9.3°. For the remaining participants (S08 and S09), retinotopic mapping stimuli were bars composed of randomly generated moving dots, which participants covertly attended to while performing a motion discrimination task (see Mackey et al., 2017 for details).

After defining retinotopic ROIs using these methods, we further thresholded the ROIs using an independent localizer task to identify voxels that were responsive to the region of space in which shape stimuli could appear (see *Silhouette localizer task* for details on this task). The data from the localizer were analyzed using a general linear model (GLM) implemented in FSL’s FMRI Expert Analysis Tool (FEAT; version 6.00). This analysis included performing brain extraction and pre-whitening (Smith, 2002; Woolrich et al., 2001). We generated predicted BOLD responses by convolving each stimulus onset with a canonical gamma hemodynamic response (phase = 0s, s.d. = 3s, lag = 6s), and combined individual runs using a standard weighted fixed effects analysis. We identified voxels that were significantly activated by the stimulus versus baseline (p < 0.05, false discovery rate (FDR) corrected). This mask of responsive voxels was then intersected with each ROI definition to obtain the final thresholded ROI definitions. The exception to this was the IPS ROIs, to which we did not apply any additional thresholding; this was because the localizer yielded few responsive voxels in IPS for some participants. See Supplementary Table 1 for the final number of voxels in each ROI, after thresholding.

### Shape stimuli

We used a set of shape silhouette stimuli that varied parametrically along two continuous dimensions, generating a 2-dimensional shape space (Figure 1A). Each shape in this space was a closed contour composed of radial frequency components (RFCs; Op de Beeck et al., 2001; Zahn & Roskies, 1972). Each shape was composed of 7 different RFCs, where each component has a frequency, amplitude, and phase. We selected these stimuli because they can be represented in a low-dimensional grid-like coordinate system, but are more complex and abstract relative to simpler stimuli such as oriented gratings. Importantly, the changes along each axis in the shape space involve variability in multiple regions of the image, so categorizing the shapes correctly required participants to integrate information globally across the image, rather than focusing on a single part of the shape. To generate the 2-dimensional shape space, we parametrically varied the amplitude of two RFCs, leaving the others constant. The manipulation of RFC amplitude was used to define an x/y grid in arbitrary units that spanned positions between 0-5 a.u., with adjacent grid positions spaced by 0.1 a.u. All shape space positions on all trials were sampled from this grid of shape space positions. We also defined a coarser grid of 16 points (a 4×4 grid) which was used to generate the 16 stimuli that were shown on the majority of trials; this grid is referred to as the “main grid”, and included all x/y combinations of the points [0.1, 1.7, 3.3, 4.9] in shape space coordinates. Stimuli corresponding to points in shape space that were not part of the main grid were used to make the tasks more difficult, see *Main task design* for details.

We divided the shape space into four quadrants by imposing boundaries at the center position of the grid (2.5 a.u.) in each dimension. To define the binary categories that were relevant for each task (see *Main task design)*, we grouped together two quadrants at a time, with the *Linear-1* task and *Linear-2* tasks grouping quadrants that were adjacent (creating either a vertical or horizontal linear boundary in shape space), and the *Nonlinear* task grouping quadrants that were non-adjacent (creating a non-linear boundary). During task training as well as before each scanning run, we utilized a “prototype” image for each shape space quadrant as a way of reminding participants of the current categorization rule. The prototype for each quadrant was positioned directly in the middle of the four main grid positions corresponding to that quadrant (i.e. the x/y coordinates for the prototypes were combinations of [0.9, 4.1] a.u.). These prototype images were never shown during the categorization task trials, to prevent participants from simply memorizing the prototypes. Shapes used in the task were also never positioned exactly on any quadrant boundary in order to prevent any ambiguity about category.

### Display parameters

During all scanning runs, stimuli were presented to participants by projecting onto a screen that was mounted on the inside of the scanner bore, just above the participant’s chest. The screen was visible to the participant via a mirror that was attached to the head coil. The image projected onto the screen was a rectangle with maximum horizontal eccentricity of 13 degrees (center-to-edge distance) and maximum vertical eccentricity of 10 degrees. In the main task and silhouette localizer task, the region of the screen in which shapes could appear subtended a maximum eccentricity of 11 degrees in the horizontal direction, and 9 degrees in the vertical direction. The fixation point in all tasks was a gray square 0.2 degrees in diameter; participants were instructed to maintain fixation on this point throughout all experimental runs.

In the main task, shapes were displayed as gray silhouettes on a gray background. For all participants except for the first participant (S01), the shapes were darker than the background (shape = 31, background = 50; luminance values are in the range 0-255). For S01, the shapes were lighter than the background (shape = 230, background = 77). The change in parameters was made because the brighter stimuli shown to S01 led to display artifacts when scanning subsequent participants, and darker stimuli reduced these artifacts. S01 reported no artifacts and performed well on the task. No gamma correction was performed.

### Main task design

The main experimental task consisted of categorizing shape silhouette stimuli (Figure 1) into binary categories. There were three task conditions: *Linear-1, Linear-2,* and *Nonlinear*, each of which corresponded to a different binary categorization rule. Shape stimuli were drawn from a two-dimensional shape space coordinate system (see *Shape stimuli*). The *Linear-1* and *Linear-2* tasks used a boundary that was linear in this shape space, while the *Nonlinear* task used a boundary that was non-linear in this shape space (requiring participants to group non-adjacent quadrants into a single category, see Figure 1 for illustration). Each trial consisted of the presentation of one shape for 1s, and trials were separated by an inter-trial interval (ITI) that was variable in length, uniformly sampled from the interval 1-5s. Participants responded on each trial with a button press (right index or middle finger) to indicate which binary category the currently viewed shape fell into; the mapping between category and response was counter-balanced within each scanning session. Participants were allowed to make a response anytime within the window of 2s from stimulus onset. Feedback was given at the end of each run, and included the participant’s overall accuracy, as well as their accuracy broken down into “easy” and “hard” trials (see next paragraph for description of hard trials), and the number of trials on which they failed to respond. No feedback was given after individual trials.

Each run in the task consisted of 48 trials and lasted 261s (327 TRs). Of the 48 trials, 32 of these used shapes that were sampled from a grid of 16 points evenly spaced within shape space (“main grid”, see *Shape stimuli*), each repeated twice. These 16 shapes were presented twice per run regardless of task condition. The remaining 16 trials (referred to as “hard” trials) used shapes that were variable depending on the current task condition and the difficulty level set by the experimenter. The purpose of these trials was to allow the difficulty level to be controlled by the experimenter so that task accuracy could be equalized across all task conditions, and prevent any single task from being trivially easy for each participant. For each run of each task, the experimenter selected a difficulty level between 1-13, with each level corresponding to a particular bin of distances from the active categorization boundary (higher difficulty denotes closer distance to boundary). These difficulty levels were adjusted on each run during the session by the experimenter, based on performance on the previous run, with the goal of keeping the participant accuracy values within a stable range for all tasks (target range was around 80% accuracy). For the *Nonlinear* task, the distance was computed as a linear distance to the nearest boundary. The “hard” trials were generated by randomly sampling 16 shapes from the specified distance bin, with the constraint that 4 of the shapes had to come from each of the four quadrants in shape space. This manipulation ensured that responses were balanced across categories within each run. For many of the analyses presented here, we excluded these hard trials, focusing only on the “main grid” trials where the same images were shown across all task conditions.

Participants performed 12 runs of the main task within each scanning session, for a total of 36 runs across all 3 sessions (with the exception of one participant (S06) for whom 3 runs are missing due to a technical error). The 12 runs in each session were divided into 6 total “parts” where each part consisted of a pair of 2 runs having the same task condition and the same response mapping (3 conditions x 2 response mappings = 6 parts). Each part was preceded by a short training run, which consisted of 5 trials, each trial consisting of a shape drawn from the main grid. The scanner was not on during these training runs, and the purpose of these was to remind the participant of both the currently active task and the response mapping before they began performing the task runs for that part. The order in which the 6 parts were shown was counter-balanced across sessions. Before each scan run began, the participant was again reminded of the current task and response mapping via a display that presented four prototype shapes, one for each shape space quadrant (see *Shape stimuli* for details on prototype shapes). The prototypes were arranged with two to the left of fixation and two to the right of fixation, and the participant was instructed that the two leftmost shapes corresponded to the index finger button and the two rightmost shapes corresponded to the middle finger button. This display of prototype shapes was also used during the training runs to provide feedback after each trial: after each training trial, the four prototype shapes were shown, and the two prototypes corresponding to the correct category were outlined in green, with accompanying text that indicated whether the participant’s response was correct or incorrect. This feedback display was not shown during the actual task runs.

Before the scan sessions began, participants were trained to perform the shape categorization tasks in a separate behavioral session (training session took place on average 4.0 days before the first scan session). During this behavioral training session, participants performed the same task that they performed in the scanner, including 12 main task runs (2 runs for each combination of condition and response mapping; i.e., each of the 6 parts). As in the scan sessions, each part was preceded by training runs that consisted of 5 trials, each accompanied by feedback. Participants completed between 1-3 training runs before starting each part. Average training session accuracy was 0.81 ± 0.02 (mean ± SEM across 10 participants) for the *Linear-1* task, 0.81 ± 0.02 for the *Linear-2* task, and 0.78 ± 0.02 for the *Nonlinear* task.

### Silhouette localizer task

A silhouette localizer task was used to identify voxels that were responsive to all the regions of retinotopic space where the shape stimuli could appear. For this task, a single silhouette shape was generated that covered the area spanned by any shape in the main grid. The silhouette region was rendered with a black-and-white flashing checkerboard (spatial period = 2 degrees) against a mid-gray background. On each trial, the flashing checkerboard silhouette stimulus appeared for a total duration of 7s, with trials separated by an ITI that varied between 2-8s (uniformly sampled). During each trial the checkerboard was flashed with a frequency of 5 Hz (1 cycle = on for 100 ms, off for 100 ms). On each cycle, the checkerboard was re-drawn with a randomized phase. There were 20 trials per run of this task, and participants performed between 4 and 7 runs of this task across all sessions. During all runs of this task, participants were instructed to monitor for a contrast dimming event and press a button when the dimming occurred. Dimming events occurred with a probability of 0.10 on each frame, and were separated by a minimum of 4 cycles. There were on average 17 dimming events in each run (minimum 10; maximum 25). Average hit rate (proportion of events correctly detected) was 0.69 ± 0.07 (mean ± SEM across 10 participants), and the average number of false alarms per run was 3.42 ± 1.41 (mean ± SEM across 10 participants).

### Image similarity analysis

To estimate the perceptual discriminability of our shape categories, we used two computer vision models to extract activations in response to each stimulus image. We first used the GIST model (Oliva & Torralba, 2001), which is based on Gabor filters and captures low-level spectral image properties. We also extracted features from a pre-trained SimCLR model (T. Chen et al., 2020), which is a self-supervised model trained using contrastive learning on a large image database. We selected these two models because the GIST model captures clearly defined image properties similar to those represented in the early visual system, while the SimCLR model can capture a wider set of image features, including mid-level and high-level properties. The GIST model was implemented in Matlab, using a 4×4 spatial grid, 4 spatial scales, and 4 orientations per spatial scale. The version of SimCLR that we used was implemented in PyTorch and used a ResNet-50 backbone (pre-trained model downloaded from https://pypi.org/project/simclr/). We extracted activations from blocks [2,6,12,15] and performed a max-pooling operation (kernel size = 4, stride = 4) to reduce the size of activations from each block. We used principal components analysis (PCA) to further reduce the size of activations, retaining a maximum of 500 components per block, and concatenated the resulting features across all blocks.

Using these activations, we computed the separability of shape categories across each of our boundaries (*Linear-1, Linear-2, Nonlinear*) by computing all pairwise Euclidean distances between main grid shapes in the same category (within-category distances) and main grid shapes in different categories (between-category distances). We then computed the average of the within-category distances (w) and between-category distances (b). The separability measure for each boundary was computed as: (b-w)/(b+w).

### Multivariate classifier analysis

We used a multivariate classifier to estimate how well the voxel activation patterns from each ROI could be used to discriminate different shape stimuli. We performed three different types of binary classification (*Linear-1, Linear-2, Nonlinear)*, as well as 16-way multinomial classification, and the following details apply to all classifier types. Classification was performed within each participant, each ROI, and each task condition separately. Before training the classifier, we mean-centered the activation patterns on each trial, by subtracting the average signal across voxels from each trial. We cross-validated the classifier by leaving one run out at a time during training, looping over held-out test runs so that every run served as the test run once. During training of the classifier, we used only trials on which main grid shapes were shown. For the 16-way classifier, we treated each of the 16 unique shapes as distinct classes. For the binary classifiers, we split the 16 shapes into two classes according to either the *Linear-1* category boundary, the *Linear-2* category boundary, or the *Nonlinear* category boundary.

Using these class labels, we then constructed a logistic regression classifier, implemented using *scikit-learn* (version 1.0.2) in Python 3.6. We used the ‘lbfgs’ solver and L2 regularization. To select the L2 regularization parameter (C), we created a grid of 20 candidate C values that were logarithmically spaced between 10^-9^ and 1. We then used nested cross-validation on the training data only to select the C resulting in highest accuracy across folds, and re-fit the model for the entire training set using the best C parameter. The resulting classifier was then used to predict the class (1-2, or 1-16) for all trials in the test dataset (note that this included trials where the viewed shape was not in the main grid, and thus was not included in classifier training). In addition to a predicted class for each trial, the classifier returned a continuous probability estimate for each of the classes, obtained using a softmax function.

To evaluate whether the accuracy of the classifier was significantly greater than chance, we used a permutation test. To do this, we performed 1000 iterations of training and testing the classifier, constructed in the same way as described above, using shuffled labels for the data. We always performed shuffling within a given scan run, so that the run labels were kept intact, and leave-run-out cross-validation was performed as in the original method. To make this computationally feasible, we did not perform C selection on every shuffling iteration, instead we used a fixed C value of 0.023 (for the 16-way classifier) or 0.007 (for each of the 2-way classifiers), which were approximately the median of the C values obtained across all models fit to the real data. We obtained a p-value for each individual participant, ROI, and task condition by computing the proportion of shuffle iterations on which shuffled classifier accuracy was greater than or equal to the real classifier accuracy. To obtain p-values for the participant-averaged classification accuracy for each ROI and task, we used the same procedure but first averaged the values across participants, within each shuffle iteration. All reported p-values were false-discovery-rate (FDR) corrected at q = 0.01 (Benjamini & Hochberg, 1995).

### Confusion matrix analysis

For each participant, ROI, and task, we generated a confusion matrix for the 16-way multinomial classifier. This was a 16 x 16 matrix where each row represents the set of trials on which a given shape was actually shown, and each column in the row represents the proportion of those trials that the classifier assigned into each of the 16 classes, and each row sums to 1. To compute confusion matrices we used only trials in the main grid, and only used trials on which the participant made a correct behavioral response. To quantify the alignment of confusion matrices with the representation needed to solve each task, we generated template confusion matrices for the *Linear-1* and *Linear-2* tasks, where each template matrix had 0 for pairs of stimuli that were on different sides of the boundary and 1 for pairs of stimuli that were on the same side of the boundary. We then computed the Pearson correlation coefficient between each actual confusion matrix and each template confusion matrix. Finally, we applied a Fisher z-transform to these correlation coefficient values, using the inverse hyperbolic tangent function (arctanh).

### Classifier confidence

To obtain a continuous estimate of the discriminability of shapes belonging to different binary categories, we computed a measure we term “classifier confidence”, which is based on the continuous probability estimates output by each binary or 16-way classifier. For each boundary and each individual trial, our measure of classifier confidence was computed as the difference between the total probability assigned by the classifier to the “correct” binary category for that trial [p(correct)] and the total probability assigned by the classifier to the “incorrect” binary category for that trial [p(incorrect)]. For each of the binary classifiers, it is straightforward to compute p(correct) and p(incorrect) based on the probability assigned to each binary class. For the 16-way classifier, we obtained p(correct) by summing the probability assigned to the 8 main grid shapes in the same category as the shape on the current trial (based on whichever category boundary was currently being considered), and p(incorrect) by summing the probability assigned to the 8 main grid shapes in the other category. This allowed us to compute classifier confidence from the 16-way classifier, with respect to each of the three category boundaries. Note that this measure of confidence can be computed even when the test trial shape is not part of the main grid. To interpret this measure, large positive values of confidence indicate high discriminability of shapes across a given category boundary, and large negative or zero values indicate poor discriminability.

For the analyses where confidence values are broken down by “far” and “near” trials, the far and near trials are always restricted to positions in the main grid. For the *Linear-1* and *Linear-2* tasks, there are 8 total positions counted as far and 8 counted as near. For the *Nonlinear* task, we counted the 4 corner positions as far and the 12 other positions as near. When average confidence values are reported, they are averaged over behaviorally correct trials only (unless otherwise specified).

### Bootstrap resampling procedures

When comparing classifier confidence values between correct and incorrect trials, we used bootstrap resampling to match the distribution of shape positions sampled on correct versus incorrect trials. This controls for the possibility that correct and incorrect trials had different stimulus properties; for example, harder trials would be more likely to be incorrect. The difference in stimulus properties could have, if not corrected, contributed to a difference in average confidence between correct and incorrect trials. This analysis was done using only “hard” trials (i.e., trials close to the boundary and not on the main grid), because these had the highest rate of incorrect responses. To perform resampling, for each boundary we collapsed the set of coordinates sampled on the “hard” trials onto a single axis that ran perpendicular to the boundary of interest. For the *Nonlinear* task, instead of collapsing coordinates onto a single axis, we computed the distance between each [x,y] coordinate and the nearest linear boundary, and multiplied by (+1) for coordinates in nonlinear category 1 or (−1) for coordinates in nonlinear category 2, which results in a single coordinate value that captures distance from the boundary as well as category sign. We then binned these coordinates into a set of 12 linearly-spaced bins that spanned the portion of shape space nearest the boundary (from 1.8 to 3.2 in shape space coordinates; see *Shape stimuli*). For each participant and task, we then identified a subset of these 12 bins that were sampled on both correct and incorrect trials, and were also symmetric around the categorization boundary. We then performed 1,000 iterations on which we resampled with replacement a set of approximately 100 correct trials and approximately 100 incorrect trials that each evenly sampled from all bins, and computed the average classifier confidence for this resampled set. The final confidence values for each participant reflect the average across these 1,000 bootstrapping iterations.

### Statistical analysis

To perform statistical comparisons of classifier confidence values and template correlation coefficient values (see previous sections) across ROIs and categorization tasks, we used repeated measures ANOVA tests, implemented using *statsmodels* in Python 3.6. To obtain non-parametric p-values for these tests (which are suitable to ensure that any violations of the assumptions of the parametric tests do not bias the results), we performed permutation tests where we shuffled the values within each participant 10,000 times, and computed F-statistics for each effect on the shuffled data. This resulted in a null distribution of F-values for each effect. The final p-values for each effect were based on the proportion of iterations on which the shuffled F-statistic was greater than or equal to the real F-statistic. F-statistics reported in the text reflect those obtained using the real (unshuffled) data. This procedure for obtaining non-parametric p-values is similar to previous work (e.g., Sprague & Serences, 2013; Sprague, Ester, & Serences, 2014; Ester, Sprague & Serences, 2015; Rademaker, Chunharas & Serences, 2019; Henderson et al., 2022); we also observed qualitatively similar results when using a parametric significance test as this permutation-based approach is more conservative.

To perform post-hoc tests for differences between tasks in each ROI, we used a paired t-test with permutation. For each ROI, we computed a t-statistic for the true difference between the conditions across participants, then performed 10,000 iterations where we randomly swapped the values within each participant across conditions, with 50% probability. This resulted in a null distribution of t-statistics. The final two-tailed p-value was obtained by computing the proportion of iterations on which the shuffled t-statistic was greater than or equal to the real t-statistic and the proportion of iterations on which the real t-statistic was greater than or equal to the shuffled t-statistic, taking the minimum and multiplying by 2.

## Code availability statement

All code required to reproduce our analyses is available at https://github.com/mmhenderson/shapeDim.

## Data availability

All data used in the present study will be deposited as MATLAB-formatted data in Open Science Framework.

## Acknowledgments

This work was supported by NEI R01-EY025872 to JS, NIMH Training Grant in Cognitive Neuroscience (T32-MH020002) to MH, the Swartz Foundation Fellowship for Theory in Neuroscience to NR, and the Kavli Institute for Brain and Mind Postdoctoral Award to NR. We thank Stephanie Nelli for helpful discussions during the inception of this project as well as Anna Shafer-Skelton and Julie Eitzen for their help with data collection. We would like to acknowledge Hans Op de Beeck for providing code that was used for stimulus generation.

## Author Contributions

MH and JS conceived the research. MH, JS, and NR designed, performed the research, analyzed data, and wrote the manuscript.

## Declaration of Interests

The authors declare no competing interests.

## Supplementary Material

**Supplementary Table 1.** Number of voxels in each ROI for each participant. Voxel counts are concatenated across hemispheres, and reflect the final number of voxels in each ROI, after thresholding each ROI (except for IPS) based on the results of the Silhouette Localizer task; see *Methods.* Note that the size of voxels differed for subjects S01-S07 (2 mm^3^ isotropic) and subjects S08-S10 (2.5 mm^3^ isotropic), which leads to smaller voxel counts for the last three subjects; see *Methods* for details on acquisition parameters.

|  | S01 | S02 | S03 | S04 | S05 | S06 | S07 | S08 | S09 | S10 |
| --- | --- | --- | --- | --- | --- | --- | --- | --- | --- | --- |
| V1 | 1866 | 1956 | 2216 | 2464 | 2156 | 2373 | 1520 | 669 | 476 | 422 |
| V2 | 1236 | 1834 | 1866 | 1584 | 1424 | 1570 | 1098 | 579 | 480 | 334 |
| V3 | 1109 | 1929 | 1200 | 1548 | 1430 | 1228 | 1710 | 487 | 424 | 383 |
| V3AB | 1013 | 1964 | 517 | 708 | 812 | 1256 | 1171 | 376 | 254 | 302 |
| hV4 | 277 | 641 | 484 | 1080 | 578 | 572 | 636 | 238 | 169 | 199 |
| LO1 | 369 | 331 | 462 | 465 | 352 | 465 | 768 | 307 | 102 | 251 |
| LO2 | 152 | 322 | 492 | 304 | 230 | 320 | 420 | 156 | 76 | 156 |
| IPS | 3054 | 2463 | 2336 | 2387 | 3395 | 2230 | 2761 | 1439 | 1838 | 1414 |

**Supplementary Table 2.** Results of two-way repeated measures ANOVA tests on the binary classifier accuracy values, with factors of ROI x Task, separately for the *Linear-1, Linear-2,* and *Nonlinear* classifiers (see Figure 2A-C for classifier accuracy values). All p-values were obtained using a permutation test, see *Methods* for details.

**Linear-1 classifier accuracy**
|  | F Value | Num DF | Den DF | p |
| --- | --- | --- | --- | --- |
| ROI | 70.81 | 7 | 63 | 0.0000 |
| Task | 0.83 | 2 | 18 | 0.4505 |
| ROI:Task | 2.22 | 14 | 126 | 0.0099 |

**Linear-2 classifier accuracy**
|  | F Value | Num DF | Den DF | p |
| --- | --- | --- | --- | --- |
| ROI | 87.64 | 7 | 63 | 0.0000 |
| Task | 3.17 | 2 | 18 | 0.0672 |
| ROI:Task | 0.28 | 14 | 126 | 0.9939 |

**Nonlinear classifier accuracy**
|  | F Value | Num DF | Den DF | p |
| --- | --- | --- | --- | --- |
| ROI | 53.37 | 7 | 63 | 0.0000 |
| Task | 1.65 | 2 | 18 | 0.2180 |
| ROI:Task | 0.63 | 14 | 126 | 0.8417 |

**Supplementary Table 3.** Results of three-way repeated-measures ANOVA tests on the binary classifier accuracy values for far and near trials, with factors of ROI, task and boundary (i.e., comparing *Linear-1* classifier versus *Linear-2* classifier). Classifier accuracy values are shown in Figure 3. All p-values were obtained using a permutation test, see *Methods* for details.

Far trials
|  | F Value | Num DF | Den DF | p |
| --- | --- | --- | --- | --- |
| ROI | 100.81 | 7 | 63 | 0.0000 |
| Task | 2.91 | 1 | 9 | 0.1217 |
| Boundary | 40.46 | 1 | 9 | 0.0003 |
| ROI:Task | 2.61 | 7 | 63 | 0.0205 |
| ROI:Boundary | 3.70 | 7 | 63 | 0.0016 |
| Task:Boundary | 0.35 | 1 | 9 | 0.5659 |
| ROI:Task:Boundary | 1.54 | 7 | 63 | 0.1727 |

Near trials
|  | F Value | Num DF | Den DF | p |
| --- | --- | --- | --- | --- |
| ROI | 65.53 | 7 | 63 | 0.0000 |
| Task | 5.37 | 1 | 9 | 0.0438 |
| Boundary | 9.33 | 1 | 9 | 0.0135 |
| ROI:Task | 0.46 | 7 | 63 | 0.8639 |
| ROI:Boundary | 3.48 | 7 | 63 | 0.0020 |
| Task:Boundary | 8.99 | 1 | 9 | 0.0113 |
| ROI:Task:Boundary | 1.21 | 7 | 63 | 0.3065 |

**Supplementary Figure 1.**
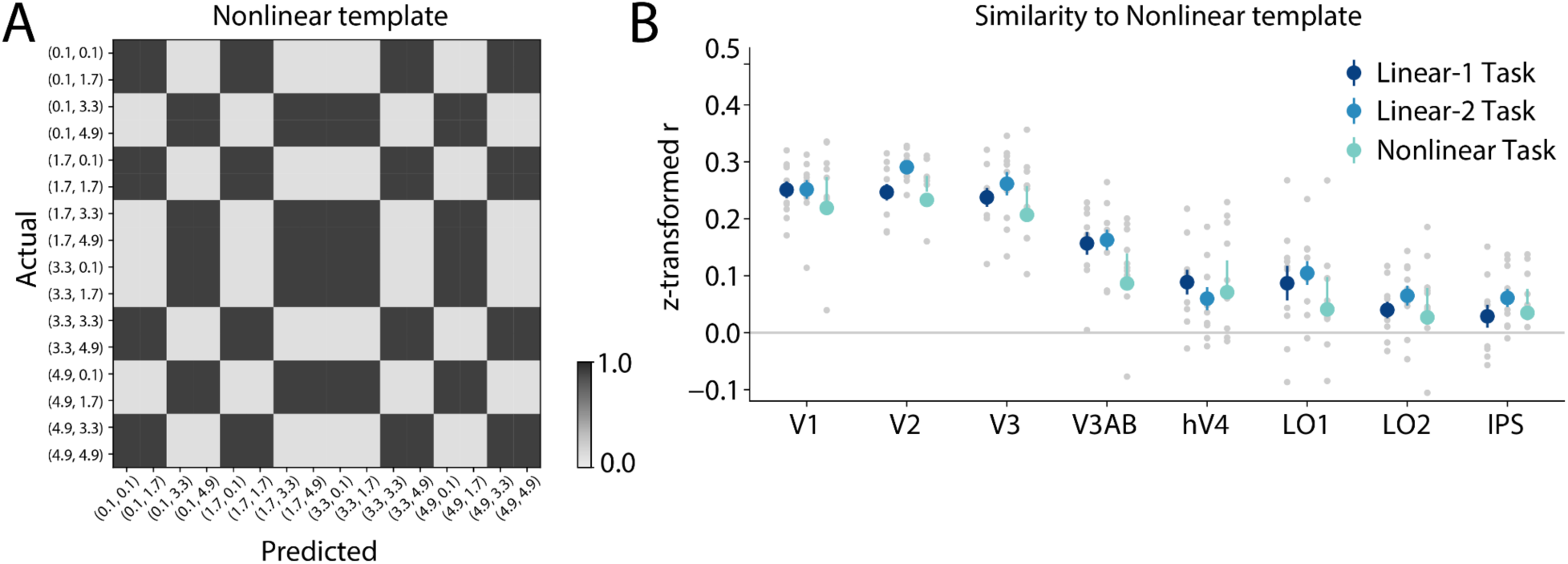
Classifier confusion matrix alignment with the *Nonlinear* template does not differ significantly across task conditions. **(A)** Template matrix for the *Nonlinear* task, representing the pattern of similarity expected for a perfect binary representation of the *Nonlinear* categorization scheme. **(B)** The similarity (Pearson correlation coefficient, z-transformed) between the *Nonlinear* template and the actual confusion matrix for each task and ROI. Gray dots represent individual participants, colored circles and error bars represent the mean ± SEM across 10 participants. A two-way repeated measures ANOVA on these similarity values revealed a main effect of ROI but no main effect of task or ROI x task interaction (ROI: F_(7,63)_ = 63.20, p < 0.001; Task: F_(2,18)_ = 1.19, p = 0.329; ROI x Task: F_(14,126)_ = 1.29, p = 0.222).

**Supplementary Table 4.** Results of three-way repeated-measures ANOVA tests on the multinomial classifier confidence values for far and near trials, with factors of ROI, task and confidence boundary (i.e., comparing *Linear-1* confidence versus *Linear-2* confidence). Classifier confidence values are shown in Figure 7. All p-values were obtained using a permutation test, see *Methods* for details.

Far trials
|  | F Value | Num DF | Den DF | p |
| --- | --- | --- | --- | --- |
| ROI | 54.44 | 7 | 63 | 0.0000 |
| Task | 0.22 | 1 | 9 | 0.6569 |
| Boundary | 49.96 | 1 | 9 | 0.0000 |
| ROI:Task | 0.82 | 7 | 63 | 0.5706 |
| ROI:Boundary | 6.58 | 7 | 63 | 0.0000 |
| Task:Boundary | 0.04 | 1 | 9 | 0.8568 |
| ROI:Task:Boundary | 1.15 | 7 | 63 | 0.3475 |

Near trials
|  | F Value | Num DF | Den DF | p |
| --- | --- | --- | --- | --- |
| ROI | 30.13 | 7 | 63 | 0.0000 |
| Task | 1.88 | 1 | 9 | 0.1975 |
| Boundary | 17.03 | 1 | 9 | 0.0011 |
| ROI:Task | 0.39 | 7 | 63 | 0.9128 |
| ROI:Boundary | 4.69 | 7 | 63 | 0.0002 |
| Task:Boundary | 11.05 | 1 | 9 | 0.0075 |
| ROI:Task:Boundary | 0.78 | 7 | 63 | 0.6281 |

**Supplementary Figure 2.**
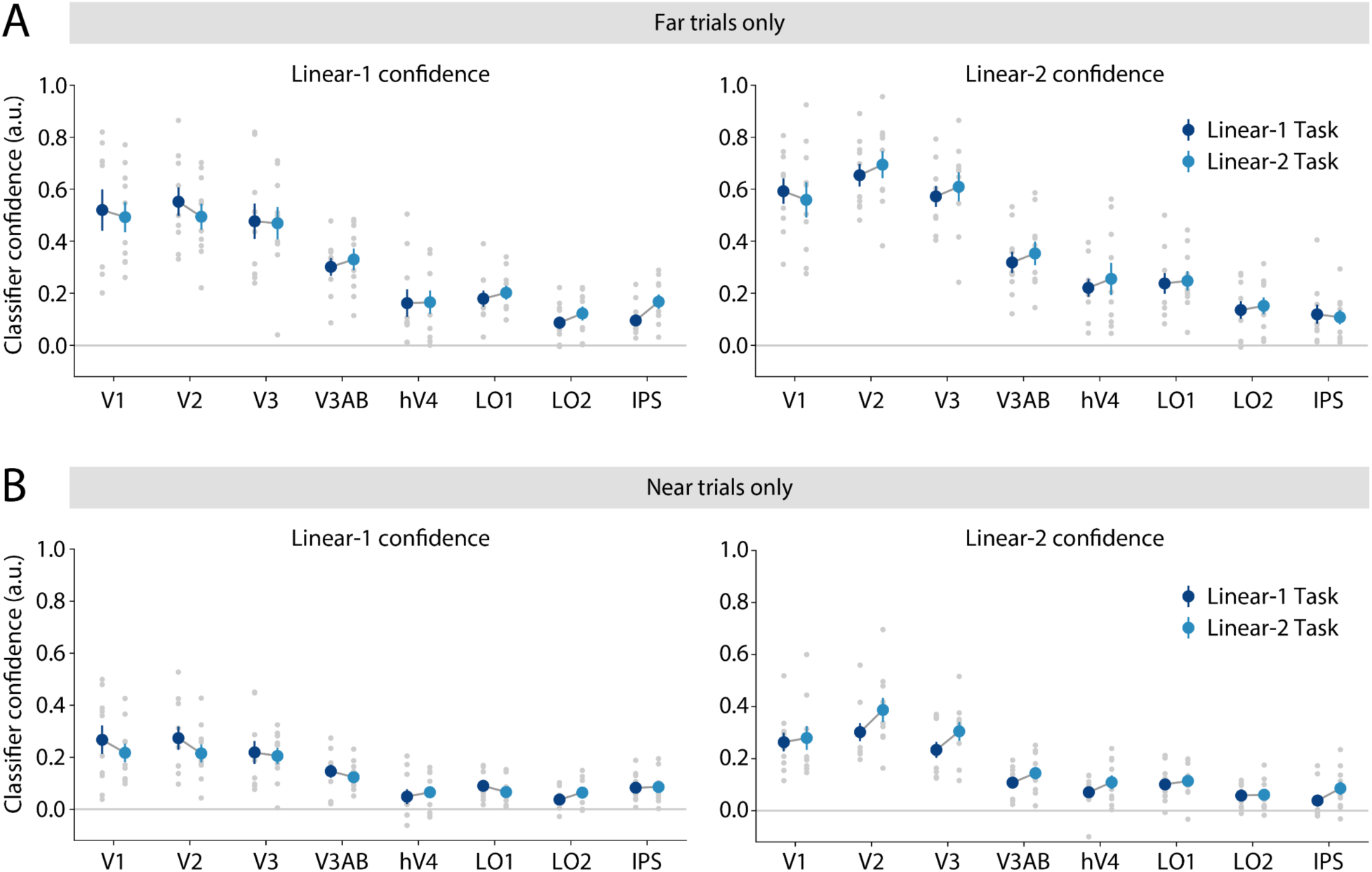
Task-related differences in classifier confidence are also measurable using binary classifiers. We used binary logistic regression classifiers that were trained to predict the category of each shape according to either the *Linear-1* or *Linear-2* decision rule (see Figure 2A-B), and computed the confidence of these classifiers for each trial as in Figure 6. **(A)** Confidence computed using “far” trials, meaning the 8 points in the main grid that fell furthest from the category boundary of interest. **(B)** Confidence computed using “near” trials, meaning the 8 points in the main grid that fell nearest to the boundary of interest. For the near trials only, we observed a main effect of task on *Linear-2* confidence (two-way repeated measures ANOVA; ROI: F_(7,63)_ = 33.81, p < 0.001; Task: F_(1,9)_ = 30.67, p < 0.001; ROI x Task: F_(7,63)_ = 0.96, p = 0.465). In (**A**-**B)**, the gray dots represent individual participants, colored circles and error bars represent the mean ± SEM across 10 participants.

**Supplementary Figure 3.**
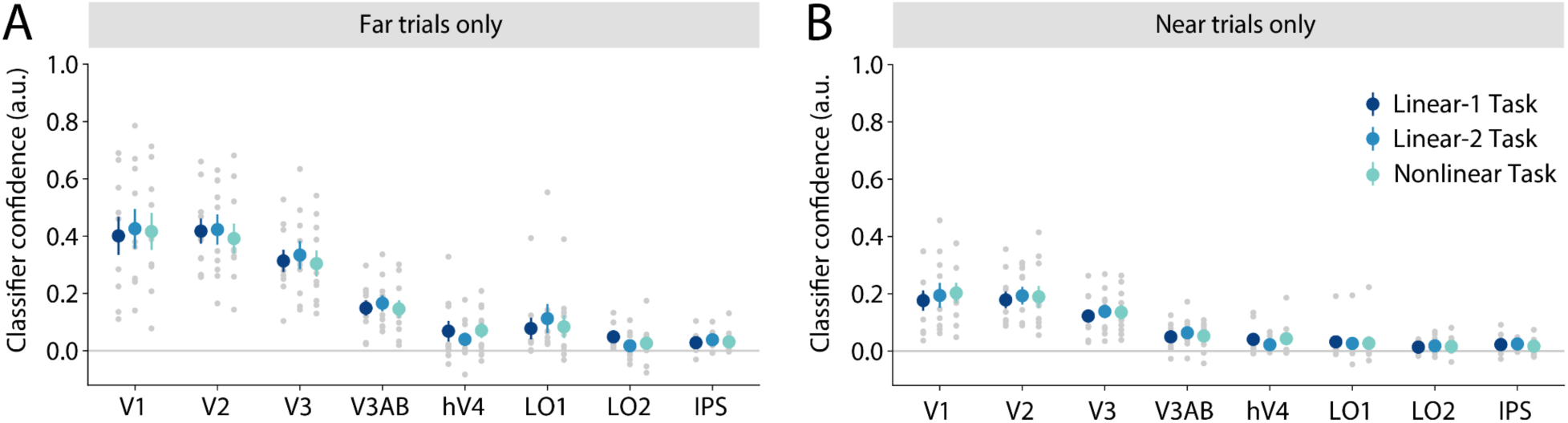
Classifier confidence across the *Nonlinear* boundary does not differ significantly across tasks. Similar to Figure 7, we computed the confidence of the classifier toward the correct *Nonlinear* task category for each trial. **(A)** Confidence computed using “far” trials, meaning the four points in the main grid that fell furthest from the two category boundaries (i.e., four corners of the shape space grid). **(B)** Confidence computed using “near” trials, meaning the 12 points in the main grid that fell nearest to either of the two category boundaries. In (**A**-**B)**, the gray dots represent individual participants, colored circles and error bars represent the mean ± SEM across 10 participants. A two-way repeated measures ANOVA on these similarity values revealed a main effect of ROI but no main effect of task or ROI x task interaction (Far trials; ROI: F_(7,63)_ = 41.70, p < 0.001; Task: F_(2,18)_ = 0.50, p = 0.618; ROI x Task: F_(14,126)_ = 0.67, p = 0.806; Near trials; ROI: F_(7,63)_ = 23.86, p < 0.001; Task: F_(2,18)_ = 0.51, p = 0.620; ROI x Task: F_(14,126)_ = 0.58, p = 0.885).

## Notes

### Competing Interest Statement

The authors have declared no competing interest.

### Summary of Updates

The manuscript has been modified to reflect the collection of three new fMRI participants, as well as the addition of new analyses.

